# Chromosome-scale reference genomes for two walnuts: *Juglans ailantifolia* and *Juglans cinerea* implicate gene presence and absence in fungal defense divergence

**DOI:** 10.64898/2026.07.29.740795

**Authors:** Cynthia N. Webster, Karl C. Fetter, Anthony He, Cristopher R. Guzman-Torres, Carolina F. Jara, Keertana Chagari, Amanda Mueller, Stefan Wnuk, Harshita Akella, Emily Strickland, Emily Trybulec, David Baukus, Laurel Humphrey, Owen McEwing, Vidya Vuruputoor, Michelle L. Neitzey, Nicole Pauloski, Aziz Ebrahimi, Emry Brannan, Martin Williams, Sean Hoban, Anna O. Conrad, James Warren, Carolyn C. Pike, Rachel J. O’Neill, Jill L. Wegrzyn

## Abstract

Butternut (*Juglans cinerea*) is a North American hardwood in rapid decline, driven largely by butternut canker disease (BCD). Within the genus, *Juglans ailantifolia* (Japanese walnut) is comparatively resistant and hybridizes readily with *J. cinerea*, yet the genomic basis of the difference is unresolved. The first chromosome-scale reference genome for *J. ailantifolia* is presented here, together with a second *J. cinerea* accession from the species’ primary range, with independent scaffolding and uniform annotation throughout. Gene family evolution was assessed across twelve Juglandaceae genomes, and a four accession pangenome contrasted the BCD resistant section Cardiocaryon (*J. ailantifolia*, *J. mandshurica*) with the BCD susceptible section Trachycaryon (*J. cinerea*, Iowa and New Brunswick accessions). Across sections separated by approximately 30 My, most defense gene families differ in presence and absence rather than copy number; five surface receptor and protease families are overrepresented in the lineage specific complement of both sections, indicating rapid bidirectional turnover. Five families differ between sections in copy number while remaining stable between the conspecific accessions, all favoring the susceptible section. The smaller set includes chitinases and NB-LRR receptors biased toward the resistant section, and dehydrins, pectin-modifying enzymes, pathogenesis-related proteins and the CBF regulon toward the susceptible one. The largest copy number difference separates the two conspecific accessions at a senescence associated cysteine protease. The northern New Brunswick accession also retains lower heterozygosity, a distinct demographic trajectory, and unique gene content enriched for calcium transport, salt stress regulation and raffinose family oligosaccharide biosynthesis, consistent with freezing tolerance.

**Significance Statement:** *Juglans cinerea* (butternut), a threatened North American walnut tree, is being lost to butternut canker disease, while *Juglans ailantifolia* (Japanese walnut) is resistant to the fungal pathogen. Chromosome-scale genomes and a pangenome place defense differences between resistant and susceptible lineages primarily in gene presence and absence. The pangenome and demographic modeling distinguish the northern *J. cinerea* accession by unique gene content consistent with cold tolerance and a distinct population history.

## INTRODUCTION

Butternut (*Juglans cinerea*) has experienced one of the steepest declines of any North American hardwood. The primary source is butternut canker disease (BCD), caused by the ascomycete *Ophiognomonia clavigignenti-juglandacearum* (*Oc-j*), first observed on *J. cinerea* in southwestern Wisconsin in 1967, formally described as *Sirococcus clavigignenti-juglandacearum* by Nair et al. (1979), and subsequently reclassified into *Ophiognomonia* (K. D. Broders and Boland, 2011). Stem girdling cankers typically kill mature trees within 5 to 10 years, and population-level mortality had already exceeded 70 to 90% across much of the native range by the late 1990s to early 2000s (K. Broders et al., 2015). Whole-genome sequencing of *Oc-j* has revealed elevated copy numbers of secreted cytochrome P450s, CAZymes, efflux pumps, and secondary metabolism gene families, providing a mechanistic basis for its capacity to overcome host defenses (Wu et al., 2020). Host resistance based on inoculation trials is thus far considered rare but there is increasing interest in characterizing individuals displaying traits associated with tolerance or resistance. Compounding pressures from another problematic fungal threat, *Armillaria* root rot, as well as habitat fragmentation, and limited natural regeneration, have accelerated its decline (LaBonte et al., 2015). As a result, *J. cinerea* is listed as endangered on the IUCN Red List and the Canadian Species at Risk Public Registry (Stritch et al. 2019; COSEWIC 2017). Although not federally listed under the United States (US) Endangered Species Act, it has been classified as critically imperiled (S1) in five states (NatureServe 2026).

Native to Canada and the United States, *J. cinerea* has a range extending from New Brunswick west to Minnesota and south to Alabama (Rink 1990). Rangewide microsatellite and chloroplast marker surveys have documented latitudinal and longitudinal gradients in diversity, reduced diversity in isolated peripheral populations (especially in the north), and chloroplast haplotype distributions consistent with multiple Quaternary refugia in the southeastern and southwestern United States, and potential independent post glacial migration (Hoban et al., 2010; Laricchia et al., 2015). More recent work by Schumacher et al. (2022) included additional northern sampling and pollen records to support the genetic distinctiveness of the New Brunswick (N.B.) *J. cinerea* population, though persistence in a northern refugium is still uncertain. A multi-year assessment conducted by Van der Meer et al. (2026) documented that the N.B. populations are undergoing rapid decline from continued *Oc-j* pressure; a decline that has already been experienced for decades throughout the rest of the range. Independent habitat suitability modeling projects a northward range shift for *J. cinerea* under changing environmental conditions, with contraction of southern habitat (Adeyemo et al., 2025). The first chromosome-scale reference assembly for the species, generated from a N.B. accession, established a baseline for ongoing genomic conservation work (Guzman-Torres et al., 2024).

*Juglans cinerea* is sympatric with *Juglans nigra* L. (black walnut) across much of its range, but no hybrid between the two has ever been confirmed (Brennan et al., 2020; Hoban et al., 2012). However, *J. cinerea* hybridizes readily with the introduced Asian *J. ailantifolia* Carrière (Japanese walnut), which is resistant to *Oc-j* and has been planted in North America since the mid-19th century (Hoban et al., 2009, 2012). The resulting hybrids, commonly known as buartnuts (*J.* × *bixbyi*), exhibit hybrid vigor and reduced canker incidence/severity in both natural populations and controlled inoculation trials (McKenna et al., 2011; Boraks and Broders, 2014; Brennan et al., 2020). Hybridization is geographically extensive, with *J. ailantifolia* almost always serving as the maternal parent (Hoban et al., 2009, 2012). As such, hybrid breeding has been considered for *J. cinerea* restoration (Pike et al., 2020), although variation in response exists within *J. cinerea,* with some families showing moderate levels of tolerance. Distinguishing *J. cinerea* from *J. ailantifolia* hybrids (buartnuts) on the landscape remains a challenge, motivating the development of nuclear and chloroplast markers (Hoban et al. 2012; Ebrahimi et al., 2025; Williams et al., 2026). Nuclear markers consistently place *J. cinerea* (the only species of section Trachycaryon) among the Asian species of section Cardiocaryon, discordant with chloroplast markers that places it sister to the New World black walnuts (Rhysocaryon) (Ebrahimi et al., 2025). This pattern is largely explained by an ancient introgression (Aradhya et al., 2007; Ebrahimi et al., 2019; Mu et al., 2020).

Chromosome-scale reference assemblies are available for representatives of all four sections of the *Juglans* genus: the widely cultivated walnut *J. regia* and its close relative *J. sigillata* (section Dioscaryon) (Marrano et al. 2020; Ning et al. 2024), the eastern North American black walnut *J. nigra* and the California endemic *J. californica* as well as the Northern California black walnut (*J. hindsii)* (section Rhysocaryon) (H. Zhou et al. 2023; Fitz-Gibbon et al. 2023), the Manchurian walnut *J. mandshurica* and Chinese walnut *J. cathayensis* (section Cardiocaryon) (Yan et al. 2021; X. Li et al. 2022; Xu et al. 2025), butternut *J. cinerea* (section Trachycaryon) (Guzman-Torres et al., 2024), and a hybrid of *J. regia* and *J. microcarpa* (Zhu et al. 2019) (Figure 1A). These resources have enabled genome-wide association studies for adaptation and agronomic traits, as well as the first walnut pangenomes linking structural variation to disease response and lipid biosynthesis (Ji et al., 2021; H. Zhou et al., 2023). Until now *J. ailantifolia,* the species most directly implicated in heritable BCD resistance and which readily hybridizes with *J. cinerea* in its native range, has lacked a reference genome.

**Figure 1.**
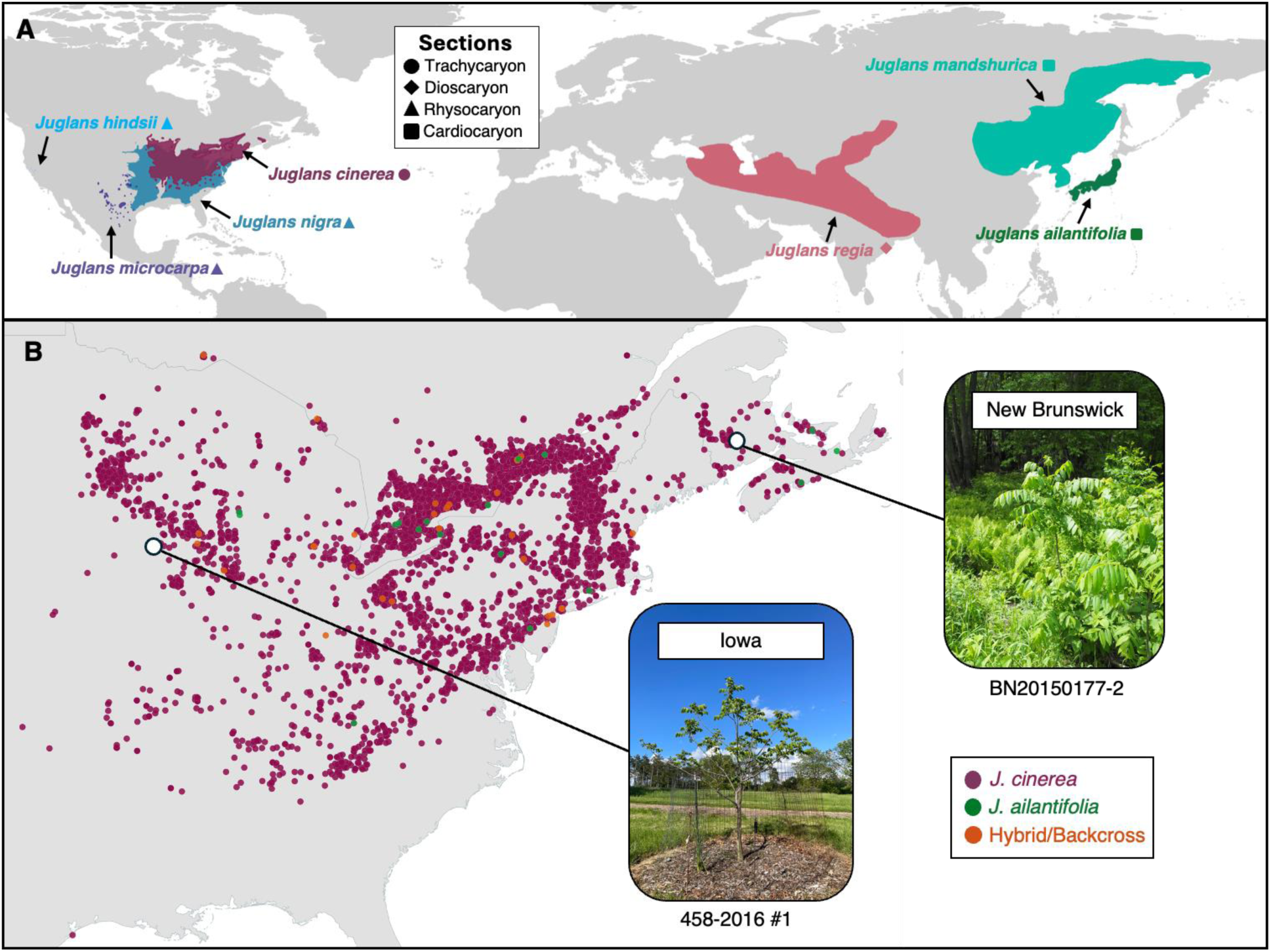
*Juglans* taxonomic ranges and North American records of *J. cinerea*, *J. ailantifolia* and their hybrids. **A)** Global distribution of *Juglans* taxa, including members from section Trachycaryon (*J. cinerea*), Dioscaryon (*J. regia*), Rhysocaryon (*J. microcarpa, J. nigra* & *J. hindsii*) and Cardiocaryon (*J. mandshurica* & *J. ailantifolia*). **B)** iNaturalist research quality grade observations of *J. cinerea* (purple), *J. ailantifolia* (green) and their hybrids and/or backcrosses (orange) across North America, along with sampling locations of the Iowa (458-2016 #1) and New Brunswick (BN20150177-2) *J. cinerea* trees used for genome assembly.

This study reports the first chromosome-scale reference assembly and annotation for *J. ailantifolia*, generated from an accession at the Morton Arboretum, together with a chromosome-scale assembly for a *J. cinerea* accession originating from the Brenton Arboretum within the species’ primary range. These complement the previously published *J. cinerea* reference from New Brunswick, Canada (Guzman-Torres et al., 2024), re-scaffolded here with independent Hi-C data to remove the bias introduced by its original scaffolding against *J. mandshurica*. All genomes, including the *J. mandshurica* assembly (X. Li et al., 2022), were annotated using a uniform informatic pipeline. These resources support a phylogenomic assessment of the *Juglans* genus, coalescent reconstruction of demographic history contrasting the *J. cinerea* accessions, comparison of repeat and methylation landscapes, confirmation of the shared juglandoid whole genome duplication, and a three species, four accession pangenome and gene family analysis characterizing how defense gene content differs between the BCD-resistant Cardiocaryon (*J. ailantifolia*, *J. mandshurica*) and susceptible Trachycaryon (*J. cinerea*) sections.

## MATERIALS & METHODS

### Plant material acquisition, extraction and sequencing

*Juglans ailantifolia* leaf tissue of accession #743-58*2 was received from the Morton Arboretum on July 11, 2023. The leaves were flash-frozen with liquid nitrogen after harvesting. The tissue was ground to a fine powder with a mortar and pestle, and stored at −80°C until extraction. Genomic DNA was extracted from the sample using a modified Vaillancourt protocol (Vaillancourt and Buell et al., 2019). Ground tissue was lysed in prewarmed Carlson Buffer (100mM Tris-HCl (pH 9.5), 2% CTAB, 1.4M NaCl, 1% PEG 8000, 20mM EDTA and 0.25% beta mercaptoethanol, 200ug/ml RNase) for 1.5 hours at 65°C. Two extractions with chloroform:isoamyl alcohol (24:1) were performed and the final upper aqueous solution was cleaned using Qiagen G-tip 100 (Qiagen, p/n 10243) following protocol. The gDNA was sheared using Covaris G-tube (pn 520079) to a size of about 25 Kb and small fragments were removed using PacBio’s Short Read Eliminator XS kit (Pacific Biosciences, pn 102-208-200). Approximately 1.5 ug of sheared and size-selected gDNA was prepared for sequencing using ONT Ligation sequencing kit (SQK-LSK114) with minor changes. DNA repair was completed at 20°C for 20 minutes followed by 65°C for 10 minutes to quench the enzyme. Elutions were performed at 37°C. The DNA was sequenced using a PromethION Oxford Nanopore Technologies (ONT) sequencer. Three loads of approximately ten fmol of the final library were run on a Promethion flow cell (ONT, FLO-PRO114M) over 96 hours. Dorado SUP accuracy model ‘dna_r10.4.1_e8.2_400bps_sup@v4.2.0’ was used to basecall reads (Wick et al., 2019; https://github.com/nanoporetech/dorado). The same protocol was followed for the *Juglans cinerea* (Iowa) accession, 458-2016 #1, received from Brenton Arboretum (Lat: 42.675, Long: −91.404) on May 15, 2024 (Figure 1B).

Snap frozen leaf tissue of *J. ailantifolia* (743-58*2) and *J. cinerea* (458-2016 #1; BN20150177-2) was sent to Arima for Hi-C sequencing using the Arima High Coverage Hi-C kit and the Arima Library Prep Module (PN A303011), according to the manufacturers protocol. Libraries were sequenced using Illumina technology in paired-end mode (150 bp) to a depth of 120 million reads per sample.

The MassARRAY SNP panel described in Williams et al. (2026) was used to validate species status, since morphological identification of pure individuals can be unreliable. Following genome assembly (see *Genome assembly* sect.), *Juglans ailantifolia* and *J. cinerea* (N.B. and Iowa) were stringently screened for sequences flanking 26 nuclear polymorphic sites using BLAST (v2.13.0), accounting for IUPAC ambiguity (Johnson et al., 2008).

### Genome assembly

*Juglans ailantifolia* and *J. cinerea* (Iowa) genome size estimates were calculated from ONT long-reads (Q>10) using kmerfreq (v.4.0) and GCE (v.1.0.2), with k-mer value 17 (Liu et al., 2013; Wang et al., 2020). The quality and quantity of the sequencing data was visualized with Nanoplot (v.1.33.0) and genome coverage was calculated from the GCE estimate (De Coster & Rademakers, 2023). Centrifuge (v.1.0.4-beta) was run against a database of fungi, bacteria, archaea and viral proteins to detect contaminants (Kim et al., 2016). Reads with a minimum hit length of 50 bp were removed, generating the final filtered read set for each genome.

Compared to Flye (v.2.9) and Hifiasm (v.0.19.6-r595), Canu (v.2.2) produced the highest quality assemblies (H. Cheng et al., 2021; Kolmogorov et al., 2019; Koren et al., 2017). Quality assessment included an estimate of contiguity with QUAST (v.5.2.0), completeness with BUSCO (v.6.0.0; Embryophyta_odbv12), and accuracy with Merqury (v.1.3) (Gurevich et al. 2013; Manni et al. 2021; Tegenfeldt et al. 2025; Rhie et al. 2020). To reduce heterozygosity, Purge Haplotigs (v.1.1.2) was applied, using minimap2 (v.2.15) for read alignment (Roach et al., 2018; H. Li, 2018).

The purged assemblies of *J. ailantifolia* and *J. cinerea* (Iowa), as well as *J. cinerea* (N.B.), from Guzman-Torres et al. (2024), were gap-filled by SAMBA (from MaSuRCA v.4.1.1) to further improve contiguity (Zimin & Salzberg, 2022). The nf-core/hic pipeline (v.2.1.0) was then used to process Illumina Hi-C reads for each genome, referencing the HindIII restriction enzyme (Servant et al., 2023; Ewels et al., 2020). The resulting HiC-Pro BAM files were fed into YaHS (v.1.2.2) to scaffold each haploid assembly over a range of resolutions (0.001-2.0 Mb) (C. Zhou et al., 2023). Juicer (v.1.2) pre and post modules produced the final scaffolded assemblies (Durand, Shamim, et al., 2016). Juicebox (v.2.17.00) was used to visualize each contact map (Durand, Robinson, et al., 2016). The expected 16 chromosomes were extracted and quality assessed (BUSCO, Merqury, and QUAST) to obtain the final metrics. Chromosomes were named using the linkage map constructed by Luo et al. (2015). Circos plots were created using shinyCircos (v2.0) (Wang et al. 2023). To generate repeat and gene density tracks, bedops (v.2.4.35) converted GFF files into BED files, and bedtools (v2.31.1) merged overlapping coordinates and calculated coverage across 10 Kb windows (Neph et al. 2012; Quinlan 2014). Methylation tracks (CpG, CHG and CHH) were generated via bedtools map (-o mean) using 100 Kb windows, and self-synteny links were created with MCScanX (v.1.0) (Wang et al. 2012).

### Genome annotation

Four genomes, excluding unplaced scaffolds, were structurally and functionally annotated with the EASEL pipeline (v.2.1.1-beta), including *J. ailantifolia, J. cinerea* (Iowa and N.B.) and *J. mandshurica* (GCA_002916435.2) (https://gitlab.com/PlantGenomicsLab/easel). For each accession, EASEL was provided with a soft-masked genome assembly (see *Repeat classification* sect.), RNA-reads, and Embryophyta OrthoDB (v.11) proteins (Kuznetsov et al., 2023). Sixty-six short, paired-end RNA-Seq libraries (>20M reads) in the *Juglans* genus were obtained from NCBI (Table S1). Fifty-six libraries were used to annotate *J. mandshurica*, and eleven were used to annotate *J. cinerea* and *J. ailantifolia* accessions. High BUSCO completeness (>95%) across StringTie2 (v2.2.3) and PsiCLASS (v.1.0.3) transcriptomes suggested that more libraries were not necessary to produce a high quality annotation (Kovaka et al., 2019; Song et al., 2019). RNA libraries were only leveraged by EASEL if mapping rates exceeded 80%. The default ‘plant’ training set was applied to each assembly. Functional information was appended to the final filtered protein sets, with and without isoforms, using EnTAP (v.2.2.0) and the RefSeq plant database (v.216) at 70/70 coverage (Goldfarb et al., 2025; Hart et al., 2020). As part of the pipeline, BUSCO (v.5.7.1) completeness (Embryophyta_odbv10) and mono:multi-ratio were calculated to evaluate each annotation. The latest version of BUSCO (v.6.0.0) was run downstream on each annotation with Embryophyta_odbv12. Metrics were also run on version 1 of *J. cinerea* (N.B.) (Guzman et al. 2024).

### Repeat classification

Lineage-specific LTR classifications were obtained using RepeatModeler (v2.0.4) with the InpactorDB plant LTR database and combined with existing RepeatModeler detected families (Flynn et al. 2020; Orozco-Arias et al. 2021). TEtrimmer (--min_blast_len 60) (v1.4.0) was run to classify unknown transposable element sequences from RepeatModeler and automate curation of transposable elements (Qian et al. 2025). In accordance with Storer et al. (2021), the top ten largest families (per genome) were manually curated. Edge extension and trimming for families were performed using the provided utilities within RepeatModeler and RepeatMasker (v4.1.5). The updated classifications from TEtrimmer and manual curation were fed into RepeatMasker to obtain repeat coverage statistics for each major transposable element type (repeatmasker.org). Subfamily classification was done using TESorter (v1.4.7) with the rexdb-plant database (R.-G. Zhang et al., 2022). Evidence of incorrect repeat classification, including missing Pfam domain annotations, prompted a more stringent TEtrimmer run (--min_blast_len 100) (v1.7.2) followed by TEsorter (v1.5.1) using updated versions. Furthermore, tandem repeat identification in *J. ailantifolia* and *J. cinerea* was conducted using TRASH (v2) to characterize putative centromeric regions (Wlodzimierz et al. 2023). HELIANO (v.1.3.1) was also run to isolate high-quality, complete Helitron annotations with the following parameters: ‘-is1 0 -is2 0 -sim_tir 90’ (Li et al. 2024).

### Gene family analysis

Gene families were inferred by OrthoFinder (v.2.5.4) using protein sequences from twelve members of the Juglandaceae family: *Juglans ailantifolia, Alfaropsis roxburghiana, Carya illinoinensis, Cyclocarya paliurus, Juglans cinerea* (N.B.)*, Juglans hindsii, Juglans mandshurica, Juglans microcarpa x regia, Juglans nigra, Juglans regia, Pterocarya stenoptera,* and *Platycarya strobilacea* (Table S2) (Emms & Kelly, 2019). Prior to characterizing genes into phylogenetic hierarchical orthogroups (HOGs), protein sequences were filtered to reflect the longest isoform using AGAT (v.1.2.0) (Dainat et al., 2022). To ensure high quality proteomes, BUSCO (v.6.0.0) was run on all the selected species (Manni et al. 2021). Original annotations sourced from NCBI were used, with the exception of *J. mandshurica*. All proteins were functionally annotated by EggNOG-mapper (v.2.1.12) to produce general descriptions and Clusters of Orthologous Genes (COG) categories (Cantalapiedra et al., 2021). Hierarchical Orthologous Groups (HOG) were assigned a function based on the longest protein sequence across all represented species. After one round of OrthoFinder, HOGs containing at least 50 sequences were removed if ≥60% of the sequences originated from a single species and the longest sequence in the cluster was annotated as a transposon, viral protein or LTR associated domain. Filtered protein sequences were re-evaluated with BUSCO to ensure completeness, and then re-assessed through OrthoFinder. OrthoFinder results were fed into Genespace (v.1.2.3) to visualize synteny across chromosome-scale assemblies (Lovell et al., 2022). *Alfaropsis roxburghiana* was used as the reference.

The ultrametric species tree and gene counts for each HOG in N0 were then passed to CAFE (v.5.1.0) to identify significantly expanding and contracting gene families (p<0.01) (Mendes et al., 2021). CafePlotter (v.0.2.0) visualized the results (https://github.com/moshi4/CafePlotter). HOGs which uniquely represented sections Cardiocaryon and Trachycaryon were extracted and functionally classified by EggNOG descriptions.

### Pangenome assembly

PanTools (v.4.3.2) was used to construct a pangenome consisting of *J. ailantifolia*, *J. mandshurica* and both *J. cinerea* accessions (Sheikhizadeh et al., 2016). The most suitable relaxation group output from the *optimal_grouping* module was 3, which set intersection rate to 0.06, similarity threshold to 75, MCL (Markov clustering) inflation to 8.4 and contrast to 6. Butternut canker disease phenotypes were not thoroughly collected for these accessions, so comparisons were made at section-level (Cardiocaryon: *J. ailantifolia* and *J. mandshurica*; Trachycaryon: *J. cinerea* (Iowa and N.B.)).

Applying a graph pangenome across two sections separated by approximately 30 My produced homology nodes that in some cases partition a single tandem array by accession rather than by gene. Physically proximal nodes were merged where their gene symbol sets intersected, so that a node with several symbols/locus identifiers could merge with a neighbour sharing any one of them. Merging was applied transitively along runs of consecutive nodes. All subsequent analyses were performed on merged units.

Because neither *J. cinerea* accession is a putative tolerant selection, the differences between them were used as a reference distribution for the variation expected between conspecific individuals. For every homology unit the within-species difference was calculated as the absolute difference in copy number between the Iowa and N.B. accessions, and the between-section difference as the difference between mean Cardiocaryon and mean Trachycaryon copy number. The 99th percentile of the within-species distribution, equal to two copies, was adopted as the threshold. Gene families were assembled from merged units; copy number was summed across the units assigned to each family and evaluated against this threshold. Presence and absence was analyzed on the same family delimitation. A unit present in both genomes of one section and absent from both genomes of the other was designated section-specific. Families were assessed for over-representation by comparing the proportion of a family’s units that were section-specific to the genome-wide proportion.

Section-specific homology units were functionally classified by EggNOG descriptions and COG categories output by EASEL. To assess GO enrichment, genes were re-annotated by the Araport11 database using EnTAP (v.2.2.0) at 50/50 coverage (C.-Y. Cheng et al., 2017; Hart et al., 2020). Similarity search accessions of Cardiocaryon and Trachycaryon were then independently exported into ClueGO (v.2.5.10) in Cytoscape (v.3.10.3) (Bindea et al., 2009, 2013; Shannon et al., 2003). *Arabidopsis thaliana* was the model organism used to define biological process and molecular function GO terms. Term significance was determined using a hypergeometric test (enrichment, depletion, two sided). Only pathways with pV ≤ 0.1 corrected with Bonferroni step down were considered.

### Methylation

Modified bases were called from raw nanopore reads using Dorado (v.0.4.2; https://github.com/nanoporetech/dorado), which performs basecalling and per-read modification calling in a single pass. Reads were aligned to the corresponding chromosome-scale assembly with minimap2 (v.2.26), sorted and indexed with samtools (v.1.16.1) (Danecek et al., 2021), and processed with Modkit (v.0.2.5-rc2; https://github.com/nanoporetech/modkit) to generate per-site methylation calls in CpG, CHG, and CHH contexts. For each site, Modkit reports the fraction of modified bases. BED outputs for each methylation type were converted to BigWig using BedGraphToBigWig (v2.10) (Kent et al., 2010). Overlapping intervals were merged to compute mean methylation per region.

Methylation profiles were visualized with deepTools (v.3.5.6) (Ramírez et al., 2014). Methylation across gene bodies and flanking regions (+/- 5 Kb) was plotted for each genome using binSize 100. To assess methylation across repeat classes, RepeatMasker files were partitioned into class-specific files, and visualized with deepTools using the same approach. To examine the spatial relationship between repeats and genes, repeat subfamilies were extracted and converted to BigWig files, and their density was plotted relative to annotated gene regions with deepTools.

### Coalescence modeling

Re-basecalled reads of both *J. cinerea* accessions were mapped to the N.B. assembly with minimap2 “-ax lr:hq” (v.2.30). SNPs were called with Clair3 (v.2.0.1) referencing model r1041_e82_400bps_sup_v400 for N.B. and r1041_e82_400bps_sup_v420 for Iowa (Su et al., 2022). Per site coverage depth was calculated with samtools (v.1.22.1) and used to filter sites with too little (<20% median) or too much (>200% median) coverage. bcftools (v.1.9) was used to filter for passing, heterozygous and biallelic sites (Danecek et al., 2021). A masking file was generated by extracting assembly gaps (Ns), GenMap (v.1.3.0) low-mappability regions (< 1.0), and depth less than 1/3 or more than 2 times the mean depth for each accession (Pockrandt et al. 2020).

Coalescence was modeled with PSMC (v.0.6.5-r74-dirty) using parameters: -N25 -t15 -r5 -p "4+25*2+4+6" for the primary run, and -N25 -t15 -r5 -b -p "2*1+20*2+4" in bootstrapping mode (Liu and Hansen 2017). To reduce spikes in recent time, as suggested by Hilgers et al. (2025), time intervals were split ("2+2+25*2+4+6"); however, this did not significantly change the initial result and thus was not pursued further. Generation time was assumed to be 30 years and the mutation rate 2.09e-8 (Zhou et al. 2023). The main model and its replicates were visualized with a custom R script.

### Whole genome duplication

Homology relationships were inferred with wgd dmd (v2) (Chen et al., 2024) using DIAMOND (v2.1.894) for initial sequence-similarity searches, followed by gene length-based bit-score normalization and MCL to construct gene families. The nucleotide (CDS) files of *J. ailantifolia*, *J. regia, J. nigra, J. cinerea* (Iowa), *J. cinerea* (N.B), *J. mandshurica*, and *Morella rubra* were provided as input. This step recovered both within genome paralogs in *J. ailantifolia*, used as anchor points for intragenomic synteny, and orthologs between *J. ailantifolia* and each of the other species. Synonymous substitution rates (Ks) were estimated for these paralog and ortholog sets with wgd ksd. The resulting distributions were visualized with wgd viz, overlaying the *J. ailantifolia* paralog Ks distribution with the pairwise ortholog Ks distributions against each species, using the accepted species tree and *J. ailantifolia* derived anchor points to orient the comparison.

## RESULTS

### Chromosome-scale genome assemblies for three Juglans accessions across two species: J. ailantifolia and ***J. cinerea***

A total of 128.5 Gb (Q>10) of ONT long reads were generated for *J. ailantifolia*, providing 223x coverage at an estimated genome size of 575 Mb (Table S3; Table S4). Filtering for contaminants removed species of bacteria and fungi, primarily *Acinetobacter baumannii, Lactiplantibacillus plantarum* and *Clostridium botulinum* (Table S5). Post-filtering, 122.8 Gb of reads remained with 213x coverage and an N50 length of 18.97 Kb. By comparison, *J. cinerea* (Iowa) had 36.7 Gb of read data, which provided an estimated 68.1x coverage based on an initial genome size of 539 Mb. Like *J. ailantifolia*, the primary filtered contaminants included *A. baumannii* and *L. plantarum*, as well as *Clostridium tetani*. The remaining 35.6 Gb of reads yielded 66.1x coverage and the genome size estimate increased to 596 Mb.

*Juglans ailantifolia* and *J. cinerea* (Iowa) filtered reads produced Canu draft genomes with 3,071 (N50: 1.63 Mb) and 2,955 (N50: 1.45 Mb) contigs, respectively, BUSCO completeness of 99.2% each, and QV scores of 40.9 (C: 92.8%) and 43.7 (C: 94.3%) (Table 1; Table S6). Purge Haplotigs improved contiguity, yielding 373 (N50: 2.39 Mb) contigs in *J. ailantifolia* and 542 (N50: 1.92 Mb) contigs in *J. cinerea* , and decreased BUSCO completeness by 0.3% and 0.1%, respectively. Following SAMBA polishing, completeness improved by 0.1% in *J. ailantifolia,* while *J. cinerea* (Iowa) was unchanged, except for an increased duplication rate. *Juglans cinerea* (N.B.) was also polished, yielding a BUSCO score of 99.1%.

**Table 1.** Final summary statistics of chromosome-scale genome assemblies and annotations produced in this study. Values in parentheses are representative of longest isoform annotation.

|  | *Juglans ailantifolia* | *Juglans cinerea*<br>(Iowa) | *Juglans cinerea* (N.B.) |  |
| --- | --- | --- | --- | --- |
| ASSEMBLY |  |  |  |  |
| Version | 1.0 | 1.0 | 2.0 | 1.0 |
| BUSCO (Dataset: Embryophyta_odb12) |  |  |  |  |
| Complete (Single & Duplicated) (%) | 97.9 | 98.8 | 99.5 | 99.7 |
| Single (%) | 83.7 | 84.0 | 84.0 | 84.0 |
| Duplicated (%) | 14.2 | 14.8 | 15.5 | 15.7 |
| Fragmented (%) | 0.4 | 0.1 | 0.1 | 0.0 |
| Missing (%) | 1.7 | 1.1 | 0.4 | 0.2 |
| QUAST |  |  |  |  |
| Number of chromosomes | 16 | 16 | 16 | 16 |
| Largest contig (bp) | 51701116 | 51522778 | 52921917 | 53338669 |
| Total length (bp) | 503742760 | 527611645 | 532501648 | 539433174 |
| GC content (%) | 36.4 | 36.6 | 36.6 | 36.6 |
| N50 (bp) | 32337293 | 34223373 | 34783554 | 34859067 |
| L50 | 7 | 7 | 7 | 7 |
| #Ns per 100 Kb | 5.1 | 7.2 | 5.4 | 5.3 |
| MERQURY |  |  |  |  |
| QV score | 41.6 | 44.1 | 43.9 | 42.9 |
| Error | 7.0E-05 | 3.9E-05 | 4.1E-05 | 5.1E-05 |
| Completeness (%) | 85.7 | 91.0 | 91.7 | 92.2 |
| ANNOTATION |  |  |  |  |
| Number of genes | 25,040 | 25,128 | 25,336 | 30,526 |
| Number of transcripts | 46,579 | 45,595 | 45,835 | 33,275 |
| Mono-exonic/multi-exonic ratio | 0.23 (0.25) | 0.23 (0.24) | 0.23 (0.24) | 0.21 (0.21) |
| **BUSCO (Dataset: Embryophyta_odb12)** |  |  |  |  |
| Complete (Single & Duplicated) (%) | 96.6 (96.3) | 97.7 (97.4) | 98.3 (98.1) | 96.8 (96.6) |
| Single (%) | 48.8 (83.5) | 51.0 (84.7) | 50.5 (84.5) | 74.0 (83.1) |
| Duplicated (%) | 47.8 (12.8) | 46.7 (12.7) | 47.8 (13.6) | 22.8 (13.6) |
| Fragmented (%) | 0.4 (0.6) | 0.1 (0.2) | 0.2 (0.3) | 2.4 (2.4) |
| Missing (%) | 3.0 (3.0) | 2.1 (2.3) | 1.4 (1.6) | 0.8 (1.0) |
| **EnTAP** |  |  |  |  |
| RefSeq plant (v216) sequence similarity (%) | 97.4 (98.4) | 97.5 (98.3) | 97.1 (98.2) | 94.1 (93.9) |
| EggNOG (v5) gene family (%) | 99.6 (99.4) | 99.6 (99.5) | 99.5 (99.4) | 99.9 (99.9) |

Going into scaffolding, *Juglans ailantifolia* had 350 contigs (N50: 2.51 Mb), *J. cinerea* (Iowa) had 507 (N50: 2.19 Mb), and *J. cinerea* (N.B.) had 326 (N50: 3.08 Mb). QV scores were 41.5 (C: 88.3%), 44.0 (C: 92.5%) and 43.7 (C: 92.4%), respectively. Finally, following Hi-C scaffolding with YaHS, each assembly had 16 observable chromosomes in the Juicer contact map (Table S7; Figure S1), with N50s ranging between 32.3-34.7 Mb, and genome sizes between 504-533 Mb, *J. ailantifolia* being the smallest. Overall, *Juglans cinerea* (N.B.) was most complete (99.5%), followed by *J. cinerea* (Iowa) (98.8%), and *J. ailantifolia* (97.9%). QV scores slightly improved for all assemblies at 43.9 (C:91.7%), 44.1 (C:91.0%), 41.6 (C:85.7%), respectively (Table 1; Figure 2). *Juglans cinerea* (N.B.) version 1 yielded a BUSCO score of 99.7%, an N50 of 34.9 Mb, and mercury QV of 42.9 (C:92.2%). While comparable to version 2, this assembly was scaffolded to *J. mandshurica*, introducing bias due to lack of Hi-C sequencing.

**Figure 2.**
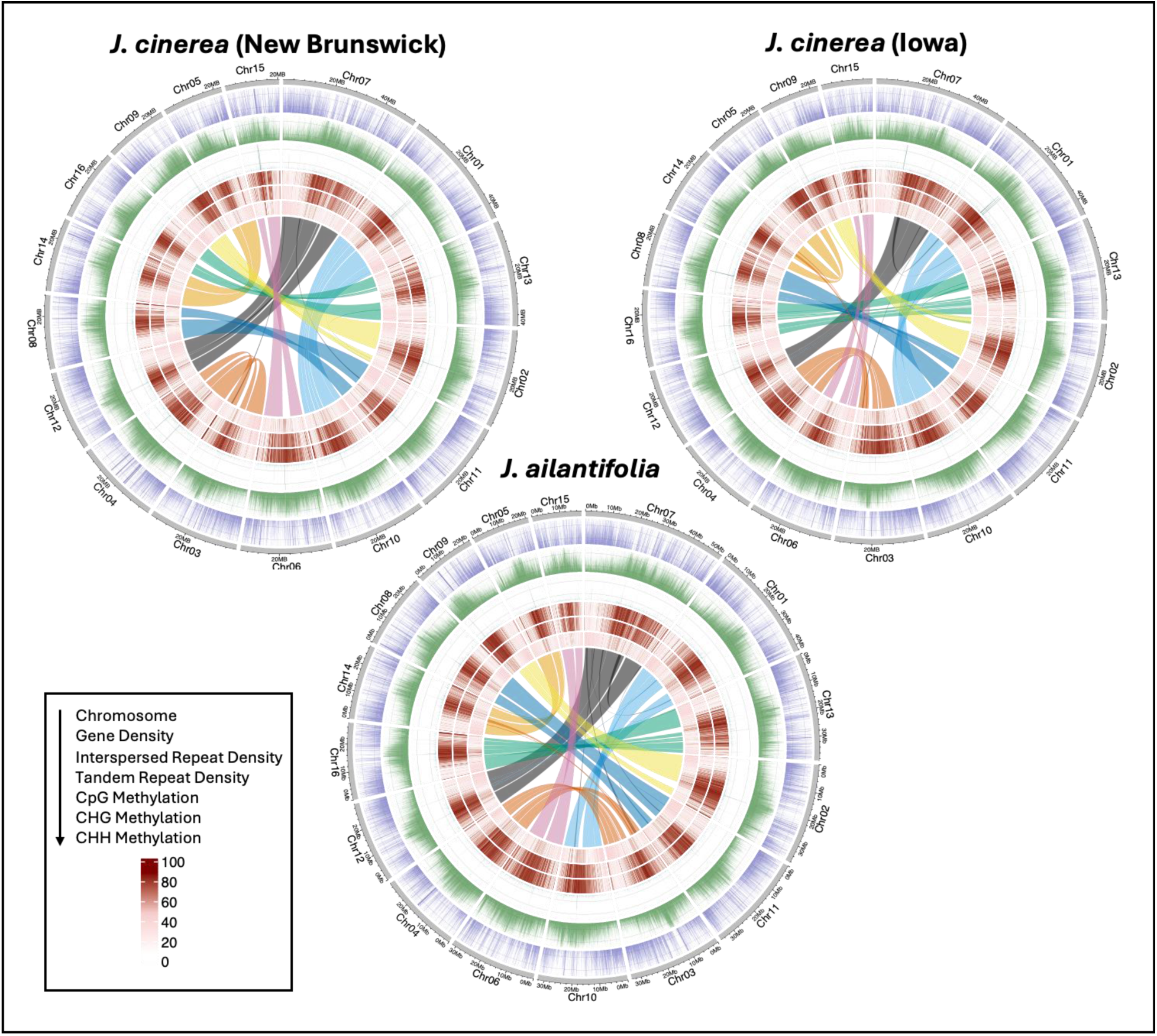
Circos plots representing chromosome-scale genome assemblies of *J. cinerea* (New Brunswick and Iowa) and *J. ailantifolia*. The tracks show (top to bottom): chromosome length (Mb), gene density (10 Kb windows), interspersed repeat density (10 Kb windows), tandem repeat density (10 Kb windows), CpG methylation (100 Kb windows), CHG methylation (100 Kb windows), and CHH methylation (100 Kb windows). The links show self-synteny, highlighting potential subgenome structure.

Across each assembly, 19/26 MassARRAY SNPs were detected based on sequence similarity to flanking sequences (Table S8). In Iowa and N.B. *J. cinerea* accessions, 89.47% of the SNPs observed were expected in *J. cinerea*, while in *J. ailantifolia* 84.21% of the SNPs observed were expected in *J. ailantifolia*.

### Protein-coding gene and transposable element annotations for four Juglans accessions across three species: J. cinerea, J. ailantifolia and J. mandshurica

The annotation process started with soft-masking repetitive elements using RepeatModeler and RepeatMasker on the chromosome-level assemblies. Going into EASEL, each genome was 43.97-44.07% masked. *Juglans mandshurica*, which was re-annotated for consistency in the pangenome analysis, was 47.02% masked (Brůna et al., 2026). RNA library mapping rates were variable across species, with *J. cinerea* (N.B.) ranging between 84.6-89.9%, *J. cinerea* (Iowa) between 82.0–88.8%, *J. ailantifolia* between 84.3-93.4% and *J. mandshurica*, which leveraged additional libraries, between 92.3-97.0% (Table S1). EASEL filtered annotations produced a conserved number of protein-coding genes across accessions. *Juglans cinerea* (N.B.) had 25,336 genes and 45,835 transcripts, *J. cinerea* (Iowa) had 25,128 genes and 45,595 transcripts, *J. ailantifolia* had 25,040 genes and 46,579 transcripts, and *J. mandshurica* had 25,687 genes and 49,910 transcripts (Table 1; Table S9). Completeness ranged between 96.7-99.2%, and mono/multi-exonic ratios ranged between 0.23-0.24, which is expected in plants (Jain et al., 2008; Vuruputoor et al., 2023).

RefSeq functional annotation rates output by EnTAP were between 97.1-97.5%. When taking the longest isoform to represent the primary gene annotation, BUSCO completeness dropped to between 96.4-98.8%, with the average duplication rate being 7.6%. This is a consequence of the longest isoform not necessarily selecting the best gene. Mono/multi-exonic ratios increased slightly, falling between 0.24-0.25. Finally, functional annotation rates increased, falling between 98.2-98.5%.

Both *J. cinerea* accessions and *J. ailantifolia* were similar by total number of repetitive elements detected, ranging from 366,493 to 385,631 total interspersed repeats over 980 to 1,071 unique repeat families. In all three accessions, 5.17%-5.84% of the genome is composed of *LINE* elements, 4.51-4.72% *DNA/TIR* elements, 14.60%-15.25% LTR elements, and 15.85%-16.30% unknown elements (Table S10). *Juglans mandshurica* had significantly higher counts at 429,882 total interspersed repeats over 1,322 unique families and showed a similar ratio between repeat types. The higher repeat count might be due to differing assembly methods as this genome was generated using PacBio sequencing instead of ONT (Li et al. 2022). Repeat analysis indicates that *LTR/Gypsy*, *LTR/Copia*, and *DNA/TIR* elements share the most recent insertion activity across all four genomes, concentrated at low divergence (0 to 5%), with *LTR/Gypsy* the most recently active of the three. *LINE* elements are the oldest class, showing a distinct peak at 35% to 45% divergence and a median Kimura distance of roughly 26% to 31%, well above the 9% to 14% range of the LTR and TIR classes (Table S11; Figure 3A). This ordering is consistent across the four genomes and with other *Juglans* assemblies.

**Figure 3.**
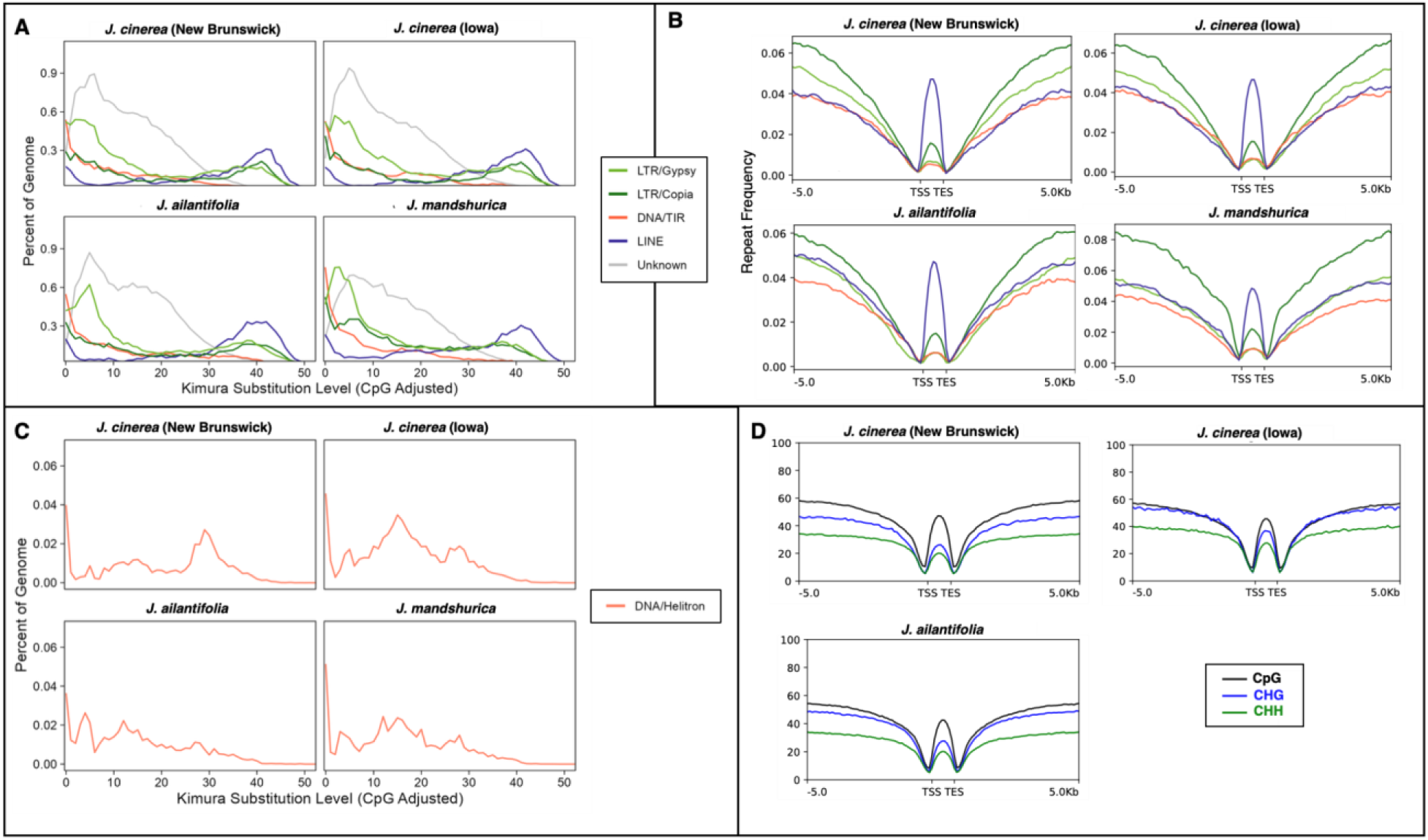
Comparative repeat and methylation landscapes across *J. cinerea* (N.B. and Iowa), *J. ailantifolia* and *J. mandshurica*. **A)** Coverage of *LTR/Gypsy, LTR/Copia, DNA/TIR, LINE* and unknown elements against Kimura divergence substitution levels. **B)** *LTR/Gypsy, LTR/Copia, DNA/TIR,* and *LINE* frequency 5 Kb up- and downstream from the translation start site (TSS) and translation end site (TES) of protein-coding genes. **C)** *DNA/Helitron* elements against Kimura divergence substitution levels. **D)** CpG, CHG , and CHH methylation patterns 5 Kb up-and downstream from the translation start site (TSS) and translation end site (TES) of protein-coding genes, excluding *J. mandshurica*.

Simple repeats accounted for ∼1.5% of detected elements in the RepeatMasker output across all four genomes. To classify high confidence tandem repeats, TRASH2 was run, resulting in estimates ranging between 0.96% and 1.19% (Table S12).

*LTR, DNA/TIR* and *LINE* repeat activity surrounding protein-coding genes showed minimal difference in insertion preferences between *J. cinerea* accessions (Figure 3B). Notably, *J. ailantifolia* and *J. mandshurica* had greater *LINE* activity up- and downstream genes relative to *J. cinerea*. Furthermore, *J. mandshurica* exhibited the highest *LTR/Copia* activity among all accessions. *LINE* elements made up the majority of intergenic TEs in every genome; across assemblies, 43% to 58% overlapped gene bodies, presenting median divergence estimates between 19% and 24%, suggesting more recent activity.

*DNA/Helitrons*, otherwise known as "peel-and-paste" transposons, represented a smaller fraction of each assembly, with values ranging between 0.44% to 0.62% in the RepeatMasker output. Their median Kimura distances ranged from 13% to 21% (Figure 3C). Additional classification with HELIANO improved annotation accuracy and reduced masking estimates to 0.14% to 0.20%, and of those elements, 78% to 89% were autonomous (Table S12). The *J. cinerea* assemblies from Iowa and N.B. contained the most *DNA/Helitrons* in their genomes, with 191 and 187 elements, respectively, compared with 145 in *J. ailantifolia* and 134 in *J. mandshurica*.

A total of 105 genes were located within +/- 5 Kb of HELIANO-classified elements in Iowa, compared with 91 in N.B., 82 in *J. mandshurica* and 78 in *J. ailantifolia* (Table S13). Both *J. cinerea* genotypes were enriched for replication protein A subunits, present as six genes in Iowa and one in N.B., and helicase-family proteins, six in Iowa, three in N.B. and one in *J. mandshurica*. Because replication initiator, helicase and replication protein A-like domains are encoded by autonomous Helitrons (Dong et al., 2011), these annotations may include internal open reading frames. Overall, 71.4% of Helitrons flanking genes were autonomous in the N.B. *J. cinerea* accession, 70.4% in Iowa, 76.4% in *J. ailantifolia* and 84.8% in *J. mandshurica* (Table S12).

### Genome-wide methylation profiling with nanopore sequencing

DNA methylation was profiled in CpG, CHG, and CHH contexts for *J. ailantifolia* and both *J. cinerea* accessions (Iowa and N.B.). *Juglans mandshurica* was obtained as a published assembly without raw ONT reads and was excluded from the methylation analysis. In plants, the three sequence contexts are maintained by largely distinct pathways, CpG primarily by MET1, CHG by the CMT3–KYP feedback loop, and CHH by RNA-directed DNA methylation and CMT2. Both CHH and CHG (non-CpG) methylation are most closely associated with the active silencing of transposable elements, particularly recently active copies near genes (Law & Jacobsen, 2010; H. Zhang et al., 2018). Each context is reported separately and describes protein coding gene bodies, gene flanking regions (UTR), and repeat bodies. The methylation profiles presented are derived from each sequenced accession and represent a single snapshot of the methylome. Thus these patterns are interpreted as methylation potential in relation to regulatory and structural features such as gene bodies, flanking promoter regions, and repeat elements.

All three genomes displayed the canonical angiosperm protein-coding gene methylation profiles, with depletion at the translation start site (TSS) and translation end site (TES) and elevated methylation in flanking regions (Figure 3D). The Iowa accession showed the highest CHH and CHG methylation across both the gene body and flanking regions, whereas the N.B. accession and *J. ailantifolia* had similar profiles across all three contexts. The most elevated signal was the non-CpG methylation of the Iowa genome; in gene bodies, the N.B. *J. cinerea* resembled *J. ailantifolia* more than it resembled its conspecific.

As expected, all repeat classes were heavily methylated across the three contexts relative to genic regions, consistent with constitutive transposon silencing (Figure S2) (Deniz et al. 2019). The elevated non-CpG methylation of the Iowa genome thus holds in both gene bodies and repeat bodies, indicating a consistent genome-wide difference. This heavy non-CpG methylation of *LTR*/*Gypsy* families in the Iowa accession coincides with the recent *LTR*/*Gypsy* expansions identified in the repeat landscape, consistent with active silencing of recently mobile elements.

### Gene family evolution across Juglandaceae

The gene family analysis with OrthoFinder, comparing twelve members of the Juglandaceae family, assigned 347,996 genes (96.4%) into 28,603 orthogroups with a mean size of 12.2 genes, and 32,756 associated HOGs (Table S14; Table S15; Table S16). The N.B. accession was used to represent *J. cinerea* in this analysis because it had the most complete proteome. Overall, 12,446 HOGs had representation from all the selected species and 2,941 were species-specific (Figure S3). Outgroup species *Pterocarya stenoptera* and *A. roxburghiana* presented the highest number of unique HOGs as expected, with 1,579 and 381, respectively. The OrthoFinder species tree placed *J. ailantifolia* sister to *J. mandshurica* within section Cardiocaryon and confirmed section Cardiocaryon as sister to section Trachycaryon (containing *J. cinerea*) (Figure 4A). Overall, chromosome-scale assemblies were highly syntenic, although orientation artifacts created apparent large inversions (Figure S4).

**Figure 4.**
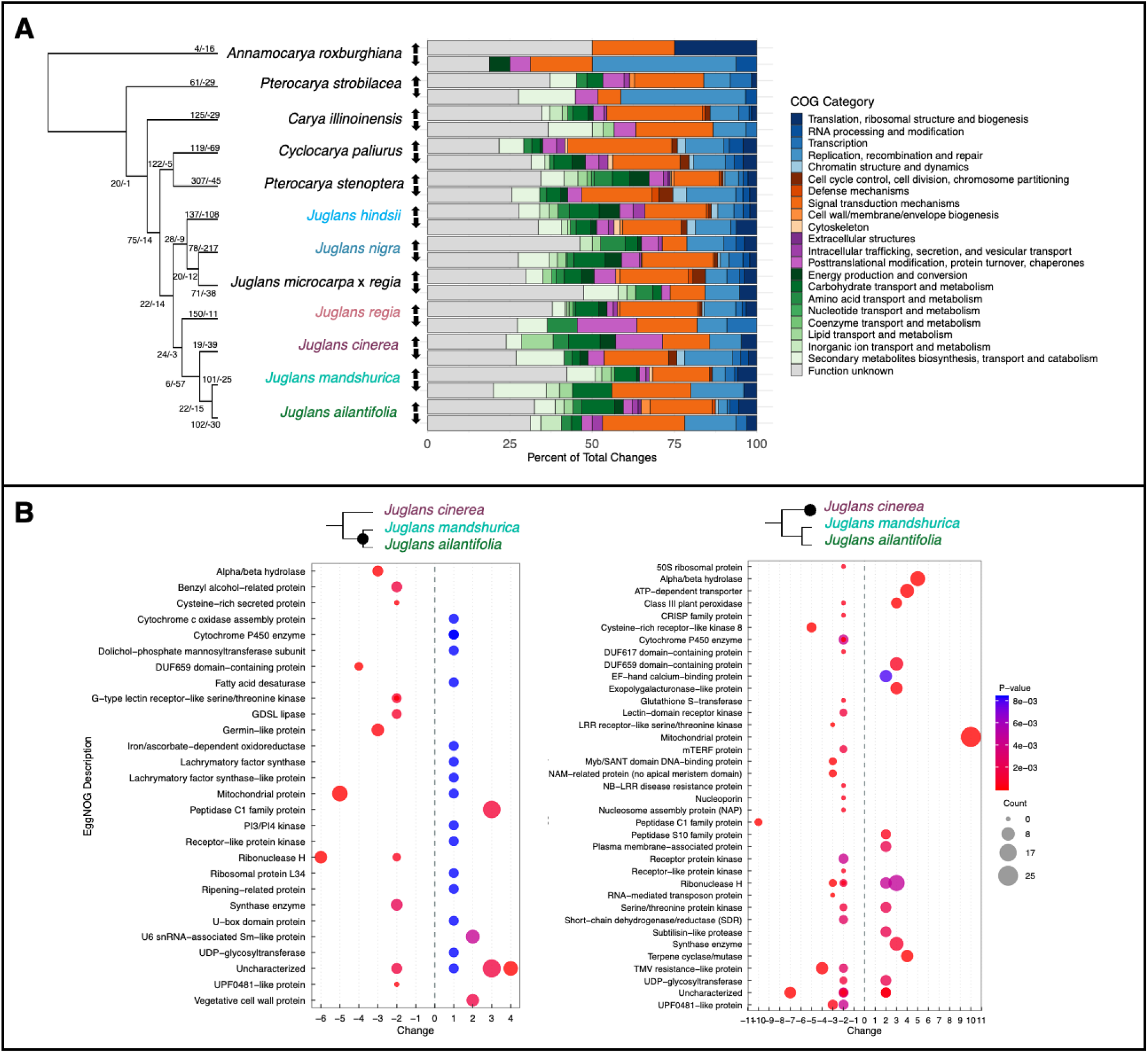
Gene family evolution in Juglandaceae. **A)** OrthoFinder phylogenetic tree with rapidly expanding and contracting gene families (p<0.01) as derived by CAFE. Percentage of total changes in expanding (up-arrow) and contracting (down-arrow) families are functionally characterized by COG category. **B)** EggNOG descriptions of ancestral node of Cardiocaryon (left) and terminal node of *J. cinerea* (right). Change is defined on the x-axis (positive = expansion; negative = contraction), p-value is described on a color gradient from dark (lower p-value) to light (higher p-value), and gene count is described by size of the circle from small to large.

Examination of gene family evolution via CAFE on the Juglandaceae protein families yielded 772 rapidly contracting and 1,613 rapidly expanding assignments across 1,145 distinct HOGs (p < 0.01) using the base model (Table S17). *Juglans nigra* had the highest number of contracting HOGs (217), while *P. stenoptera* had the highest number of expanding (307). Although many HOGs had unknown COG function (S), the most represented assigned category was signal transduction mechanisms (T) (Figure 4A).

The Cardiocaryon ancestral node yielded 22 rapidly expanding HOGs and 15 contracting (Figure 4B). Functionally annotated families with the greatest positive change included peptidase C1 family protein (+3), U6 snRNA associated Sm-like protein, vegetative cell wall protein and PI3/PI4 kinase; three cytochrome P450 HOGs also expanded, each by a single copy. Contracting families included ribonuclease H protein (−6), mitochondrial protein (−5), alpha/beta-hydrolases superfamily protein and germin-like protein subfamily 1 member (−3 each).

The *J. cinerea* (N.B.) terminal node produced 19 expanding gene families and 39 contracting (Figure 4B). Mitochondrial protein (+10), alpha/beta hydrolase (+5), ATP-dependent transporter (+4), terpene cyclase/mutase (+4) and ribonuclease H protein (+3) were amongst the top expanding, whereas peptidase C1 family protein (−10), cysteine-rich receptor-like kinase 8 (−5), TMV resistance protein N-like (−4) and LRR receptor-like serine/threonine-protein kinase (−3) were top contracting. Four cytochrome P450 HOGs also contracted at this node, each by two copies. The terminal node of *J. ailantifolia* had 102 expansions and 30 contractions, with protein receptor kinase (+4), disease resistance NB-LRR (+4), receptor protein kinase, wall associated receptor kinase-like, germin-like protein and glycosyl hydrolase 18 (+3 each) among the top expanding, and mitochondrial protein (−6), cysteine-rich receptor-like kinase 8 (−4), receptor-like protein kinase (−4) and TMV resistance protein N-like (−3) contracting. *Juglans mandshurica* had 101 expanding families and 25 contractions, with TMV resistance protein N-like (+5), protein kinase (+4), cysteine-rich receptor-like kinase 8 (+3) and chitin binding (+3) showing the greatest positive change, and protein kinase (−6), ATPase activity (−5), DDE superfamily endonuclease (−4), germin-like protein subfamily 1 member (−3) and cytochrome P450 (−3) contracting.

Several families changed in more than one terminal lineage, and their directions were not consistent with section membership. Cysteine-rich receptor-like kinase 8 (N0.HOG0000040) contracted in both *J. cinerea* (−5) and *J. ailantifolia* (−4) while expanding in *J. mandshurica* (+3), and TMV resistance protein N-like (N0.HOG0000015) followed the same pattern, contracting in *J. cinerea* (−4) and *J. ailantifolia* (−3) while expanding in *J. mandshurica* (+5). Loss within these receptor families was not restricted to Trachycaryon, the susceptible section. Peptidase C1 protein (N0.HOG0000169) expanded at the Cardiocaryon ancestral node (+3) and contracted sharply in *J. cinerea* (−10), the largest single change recovered for any defense associated family in the analysis. Germin-like protein changes involved separate HOGs in each direction, with N0.HOG0000288 contracting at the Cardiocaryon ancestral node and in *J. mandshurica* and N0.HOG0000799 expanding in *J. ailantifolia*, so the family did not follow a single trajectory.

### Pangenome assembly of section Trachycaryon and Cardiocaryon

The pangenome composed of *J. cinerea* accessions (Iowa and N.B.), *J. ailantifolia* and *J. mandshurica*, produced 17,060 core homology units with functions broadly related to transcription, RNA processing and modification, and posttranslational modification (Table S18; Table S19; Table S20; Figure 5A; Figure S5), alongside 2,592 accessory and 1,794 accession-unique units. On average, the coding sequences of core genes were longer than accessory and unique (p < 0.0001) (Figure S6). Furthermore, core genes, which are usually under high selective pressure, had the lowest repeat density 5 Kb before the translation start and after the end site, as well as within the gene body (Figure 5B).

**Figure 5.**
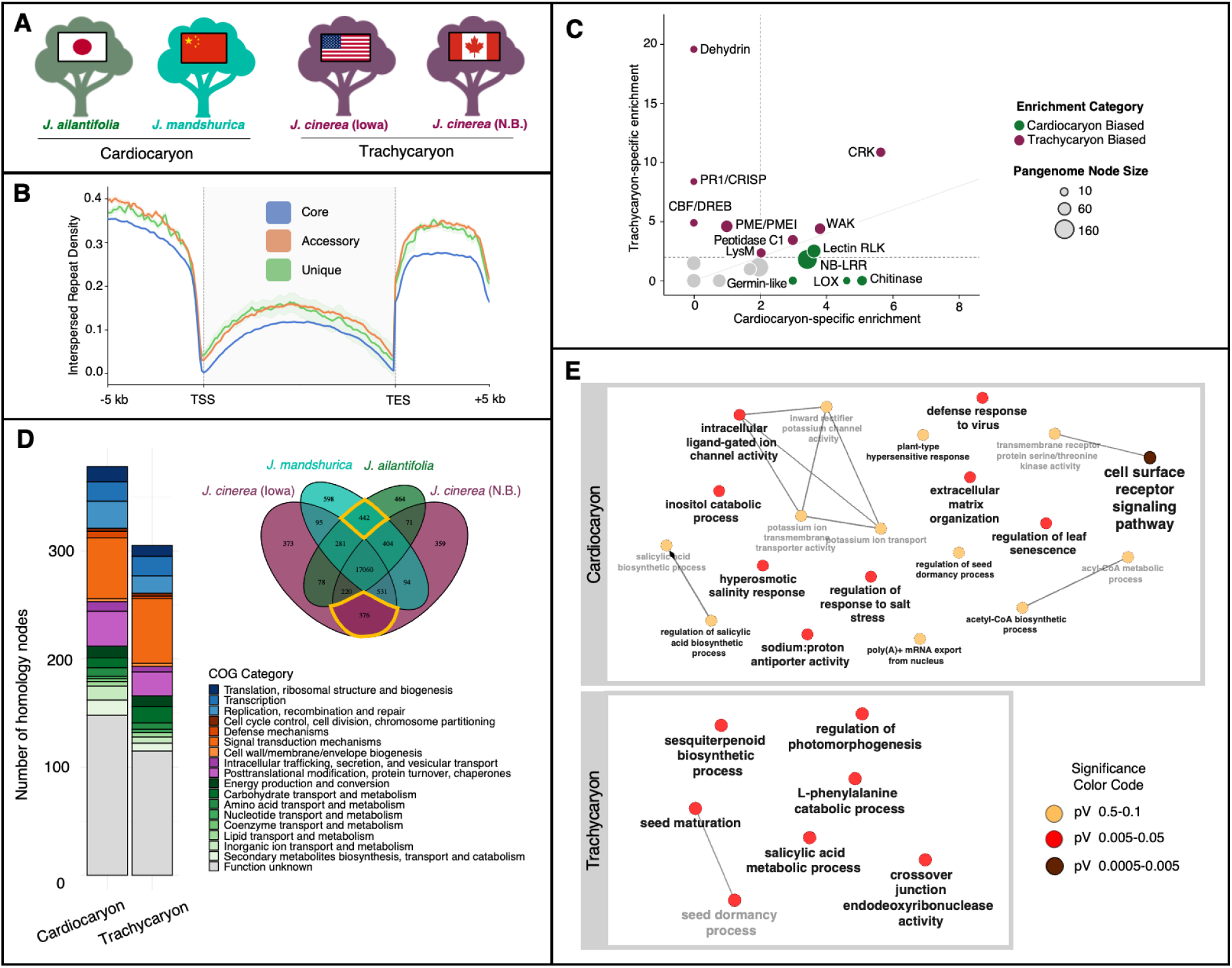
*Juglans cinerea*, *J. ailantifolia* and *J. mandshurica* pangenome assembly. **A)** Four genome accessions used to build pantools pangenome. **B)** Interspersed repeat density surrounding core (blue), accessory (orange) and unique (green) genes in pangenome 5 Kb up-and downstream from the translation start site (TSS) and translation end site (TES). **C)** Cardiocaryon vs Trachycaryon specific enrichment of homology units related to defense and stress gene families. **D)** Venn diagram showing core, dispensable and unique homology units. Circled on the venn diagram are units specific to section Cardiocaryon (top) and Trachycaryon (bottom). The stacked barplots show distribution of COG categories (Multiple COGs e.g., EG were split and counted separately, which is why there are more). **E)** GO enrichment of homology units unique to section Cardiocaryon and Trachycaryon. P-value significance is shown on a color gradient from light to dark, and the number of genes is between one and seven.

Applying a graph pangenome across two sections separated by approximately 30 My produced consecutive homology nodes assigned the same gene symbols but occupied by different accessions. Under the merge criteria, 6,165 of 27,611 homology nodes (22.3%) were absorbed into 4,582 merged homology nodes, leaving 21,446 units in total (Table S19; Table S20). Most merges joined two nodes, with the largest spanning 16. For example, ROP/RAC small GTPases occupied nodes 78381691 to 78381694, each with the same five gene symbols but holding 11 copies in Iowa, 11 in N.B., 11 in *J. mandshurica* and 10 in *J. ailantifolia*, with each node otherwise empty; merged, the unit carried 11, 11, 12 and 11 copies.

Annotation was heterogeneous within nodes, and increasingly so with node size (Table S19). Across the pangenome, 5.0% of nodes contained transcripts assigned more than one Araport11 gene name, rising from 4.2% of nodes with two to four transcripts, to 20.3% of nodes with five to nine, and 43.7% of nodes with ten or more.

### Abiotic and biotic stress-response gene copy number variation between sections

Copy number was summed across all units assigned to each of 24 defense and stress gene families (Table S21; Figure 5C). A family was called as differing between sections where the section difference exceeded the two-copy threshold and the two conspecific accessions differed by less than two copies. Five families met both criteria: pectin methylesterases and their inhibitors (65, 64, 59 and 62 copies in Iowa, N.B., *J. mandshurica* and *J. ailantifolia*), terpene synthases (22, 21, 17 and 18), subtilases (44, 45, 42 and 41), WRKY transcription factors (84, 85, 82 and 78) and dehydrins (9, 8, 6 and 6). All five differ by 2.5 to 4.5 copies and all favor Trachycaryon.

Polygalacturonases and pectate lyases have a larger section difference than any of these five, at 9.5 copies (66, 64, 56 and 55), also favoring Trachycaryon, but the two *J. cinerea* accessions differ by two copies, placing the family at the threshold rather than below it.

Six families showed section differences above two copies that failed the stability criterion, because the two conspecific accessions were as variable or more so. Chitinases and chitin binding proteins totaled 44, 51, 61 and 55 copies across 30 units, a section difference of 10.5 copies against a within-species difference of 7, and NB-LRR disease resistance proteins totaled 135, 129, 153 and 131 copies across 163 units, a section difference of 10 against a within-species difference of 6. Late embryogenesis abundant proteins (72, 79, 79 and 83) and cytochrome P450s (251, 236, 242 and 239) gave section differences of 5.5 and 3 copies against within-species differences of 7 and 15. The largest within-species differences in the dataset were at peptidase C1, including *SAG12* (56, 42, 56 and 53 copies; within-species difference 14), and germin-like proteins (59, 72, 52 and 64; within-species difference 13), both of which also gave section differences above the threshold.

Named cold-response regulators were invariant in copy number across the four accessions, whether at one copy (*CBF1*, *CBF4*, *DREB1A*, *DREB2A*, *HOS10*, *HOS15*) or at two (*CBF2*, *ICE1*, *HOS1*) (Table S20). The exception was one calmodulin binding transcription activator unit present in *J. mandshurica* alone; the remaining seven units assigned to that family were invariant.

Senescence associated gene *SAG12* was tandemly arrayed and present at high copy number in *J. mandshurica*, *J. ailantifolia* and *J. cinerea* (Iowa), but at a single copy in *J. cinerea* (N.B.) (homology unit 78381995: Iowa = 16, N.B. = 1, *J. mandshurica* = 15, *J. ailantifolia* = 13). All 45 member transcripts carried the same gene symbol and the unit comprised a single node. In Iowa the copies formed a single tandem array of 15, whereas in *J. mandshurica* they were distributed across three arrays of seven, five and three, and in *J. ailantifolia* across three arrays of seven, three and three (Table S22). A second *SAG12* unit (78403776) carries copies in the two Cardiocaryon genomes alone and is reported among the section-specific units below.

### Functional enrichment describing section-specific gene families

Isolating homology units specific to each section, Cardiocaryon had 442 and Trachycaryon 376 (Table S23; Table S24; Figure 5D). Excluding transposon-derived units, 412 units were Cardiocaryon-specific and 356 Trachycaryon-specific, representing 2.0% and 1.7% of non-transposon units respectively. Of the 24 curated families, five were overrepresented more than twofold among section-specific units in both sections: wall associated kinases (3.8-fold in Cardiocaryon, 4.4-fold in Trachycaryon), cysteine-rich receptor-like kinases (5.6 and 10.9), peptidase C1 proteins (3.0 and 3.5), LysM-domain proteins (2.0 and 2.3) and lectin-domain receptor kinases (3.6 and 2.5). Four were overrepresented in Cardiocaryon alone: chitinases and chitin binding proteins (5.1-fold, 3 units), lipoxygenases (4.6-fold, 1 unit), NB-LRR disease resistance proteins (3.4-fold, 11 units) and germin-like proteins (3.0-fold, 1 unit). Four were overrepresented in Trachycaryon alone: dehydrins (19.6-fold, 3 units), pathogenesis-related CRISP proteins (8.4-fold, 1 unit), the *CBF*/*DREB* cold regulon (4.9-fold, 1 unit of the 12 assigned to the family) and pectin methylesterases and their inhibitors (4.6-fold, 4 units). Dehydrins represent the strongest section-specific enrichment recovered for any curated family. The remaining eleven families were not overrepresented in either section, and seven of those also showed no section difference in copy number above the threshold (Table S21).

Among the genes restricted to Cardiocaryon were non-specific disease resistance 1 (*NDR1*) and Resistance to *Peronospora parasitica* protein 13 (*RPP13*), which in *A. thaliana* and other angiosperms are necessary for a hypersensitive immune response, senescence associated gene 12 (*SAG12*), a senescence-specific cysteine protease, and lysm domain GPI-anchored protein 2 precursor (*LYM2*), a chitin elicitor binding protein (Faulkner et al., 2013; James et al., 2018; M. Li et al., 2022; Samaradivakara et al., 2022; B. Yuan et al., 2024). The set also included senescence associated carboxylesterase 101 (*SAG101*), an EDS1 family protein required for TIR-domain receptor signaling, and the wall associated kinase-like gene *WAKL7*. Wall associated kinases are increasingly recognized as a central component of plant immunity, with paralogs functioning as cell-wall-localized pattern-recognition receptors that detect oligogalacturonide damage signals and pathogen effectors and trigger downstream defense signaling (Giovannetti & Genre, 2024; Pollegioni et al., 2012; Stephens et al., 2022). Consistent with turnover in this family occurring in both lineages, *WAK2* occupied three section-specific units, two restricted to Cardiocaryon and one to Trachycaryon. The Trachycaryon-specific set included the salicylic acid methylesterase *MES3*.

Cardiocaryon homology units had enriched (p < 0.1) biological process GO terms related to sodium:proton antiporter activity (GO:0015385), poly(A)+ mRNA export from nucleus (GO:0016973), plant-type hypersensitive response (GO:0009626), inositol catabolic process (GO:0019310), extracellular matrix organization (GO:0030198), hyperosmotic salinity response (GO:0042538), defense response to virus (GO:0051607), regulation of leaf senescence (GO:1900055), regulation of response to salt stress (GO:1901000), regulation of seed dormancy process (GO:2000033), acyl-CoA metabolic process (GO:0006637), acetyl-CoA biosynthetic process (GO:0006085), cell surface receptor signaling pathway (GO:0007166), transmembrane receptor protein serine/threonine kinase activity (GO:0004675), salicylic acid biosynthetic process (GO:0009697), regulation of salicylic acid biosynthetic process (GO:0080142), inward rectifier potassium channel activity (GO:0005242), potassium ion transport (GO:0006813), potassium ion transmembrane transporter activity (GO:0015079), and intracellular ligand-gated ion channel activity (GO:0005217) (Figure 5E; Table S25). Molecular function enrichment included RNA polymerase II complex binding (GO:0000993), ATP citrate synthase activity (GO:0003878), potassium ion transmembrane transporter activity (GO:0015079), intracellular ligand-gated ion channel activity (GO:0005217), and inward rectifier potassium channel activity (GO:0005242).

In Trachycaryon, enriched biological process GO terms included L-phenylalanine catabolic process (GO:0006559), crossover junction endodeoxyribonuclease activity (GO:0008821), salicylic acid metabolic process (GO:0009696), regulation of photomorphogenesis (GO:0010099), sesquiterpenoid biosynthetic process (GO:0016106), seed dormancy process (GO:0010162), and seed maturation (GO:0010431) (Figure 5E; Table S25), while molecular function included crossover junction endodeoxyribonuclease activity (GO:0008821), (E)-beta-ocimene synthase activity (GO:0034768), and myrcene synthase activity (GO:0050551).

Several CAFE results correspond in direction to differences recovered independently in the pangenome. Chitin-directed families expanded in both Cardiocaryon terminal lineages, with chitin binding in *J. mandshurica* and glycosyl hydrolase 18 in *J. ailantifolia*, and no chitin-class family changed at the *J. cinerea* node. At the *J. cinerea* (N.B.) terminal node, terpene cyclase/mutase (+4), an exopolygalacturonase-like family (+3) and a subtilisin-like protease (+2) expanded while a lipoxygenase contracted (−2), each matching the direction of the corresponding pangenome bias (Table S17; Table S21).

### Functional enrichment describing Juglans cinerea accession-specific gene families

Across the 21,446 merged units, the Iowa and N.B. accessions differed in copy number at 13.6% of units (Table S20). The 95th percentile of this within-species distribution was one copy and the 99th percentile two copies. The *J. cinerea* Iowa accession had 373 unique homology units and the N.B. accession 359, comprising 401 and 387 gene copies respectively, against 598 unique units in *J. mandshurica* and 464 in *J. ailantifolia*. The two individuals of the same species had fewer unique units than either of the two Cardiocaryon species, as expected, though the difference was less than two-fold. The functional composition of the two sets was similar, including: receptor-like protein kinases, NB-LRR and other disease resistance proteins, cytochrome P450s and Ras-related proteins the most frequent categories in both, so the accessions differed in which members of these families they hosted.

While water channel activity (GO:0015250) overlapped both accessions, the Iowa accession was uniquely enriched for potassium ion transmembrane transporter activity (GO:0015079), organic acid transport and transmembrane transporter activity (GO:0006835, GO:0015743, GO:0046943), amino acid transport (GO:0015800, GO:0015804), and mitochondrial energy metabolism (GO:0019646, GO:0042775). Comparatively, N.B. was enriched for primary shoot apical meristem specification (GO:0010072), RNA editing and processing (GO:0016554, GO:0016891), peptidyl-serine phosphorylation (GO:0018105), nucleobase biosynthetic process (GO:0046112), water transport (GO:0006833), regulation of response to salt stress (GO:1901000), far-red light and gibberellic acid mediated signaling (GO:0010018, GO:0009740, GO:0009939), positive regulation of signal transduction (GO:0009967), raffinose family oligosaccharide biosynthetic process (GO:0010325), cellular responses to nitrogen compounds (GO:1901699, GO:1902170), and ion transport and homeostasis, including cation, potassium, calcium, and proton transport activities (GO:0005261, GO:0005217, GO:0005249, GO:0006816, GO:0019829, GO:0006874, GO:0015662, GO:0015085, GO:0005262, GO:0005388, GO:0008553, GO:0051453) (Table S26).

### Helitron associated gene organization in the pagenome

Based on pangenome homology assignments, *DNA/Helitrons* flanked genes belonging to tandem arrays in all four accessions, with nine genes in N.B., five in *J. mandshurica*, eight in Iowa and four in *J. ailantifolia*, distributed across 17 homology units of which 14 spanned more than one homology unit (Table S13; Table S22). Functional annotation identified a cytochrome P450 (*CYP72A14*, homology unit 78388831) flanking an element in both *J. cinerea* accessions, where the three-node unit had five copies in Iowa against three in each of the other accessions, and a methionine gamma-lyase (*MGL*, homology unit 78388374) flanking an element in all four accessions, where the two node unit represented three copies in every genome. Of 243 homology units containing at least one Helitron-flanking gene, 15 were flanked in all four accessions, five in three, 50 in two and 173 in a single accession (Table S13).

Two loci combined *DNA/Helitron* association with accession-specific gene organization. In the N.B. accession, two tandemly arrayed genes with the cysteine-rich receptor-like protein kinase symbols *CRK25* and *EP1* in homology unit 78395005 (M3706) uniquely flanked a hypo-methylated, 26,467 bp non-autonomous *DNA/Helitron* with domains related to serine/threonin kinase activity, salt stress response/antifungal, and antifungal protein ginkbilobin-2 (Bourdais et al., 2015) (Figure 6A; Table S13; Table S22). The *EP1* annotation, while mono-exonic and absent from other species in this unit, had a complete open reading frame and did not present evidence of pseudogenization (Xie et al., 2019). Gene copies in unit 78395005 (M3706), which also carried *CRK26* and *CRK29*, totaled 3, 6, 8 and 7 in Iowa, N.B., *J. mandshurica* and *J. ailantifolia.* Comparable tandem arrays occurred in *J. mandshurica* and *J. ailantifolia*, though none of the genes resided near *DNA/Helitrons*. Flavin-containing monooxygenase 1 (*FMO1*) at homology unit 78386245 carried an additional dispersed copy in *J. ailantifolia* upstream from a hyper-methylated, 2682 bp autonomous *DNA/Helitron* containing the DEAD-box helicase PIF1 domain and Helitron helicase-like domain at N-terminus (Iowa = 1, N.B. = 1, *J. mandshurica* = 1, *J. ailantifolia* (Figure 6B; Table S13; Table S20).

**Figure 6.**
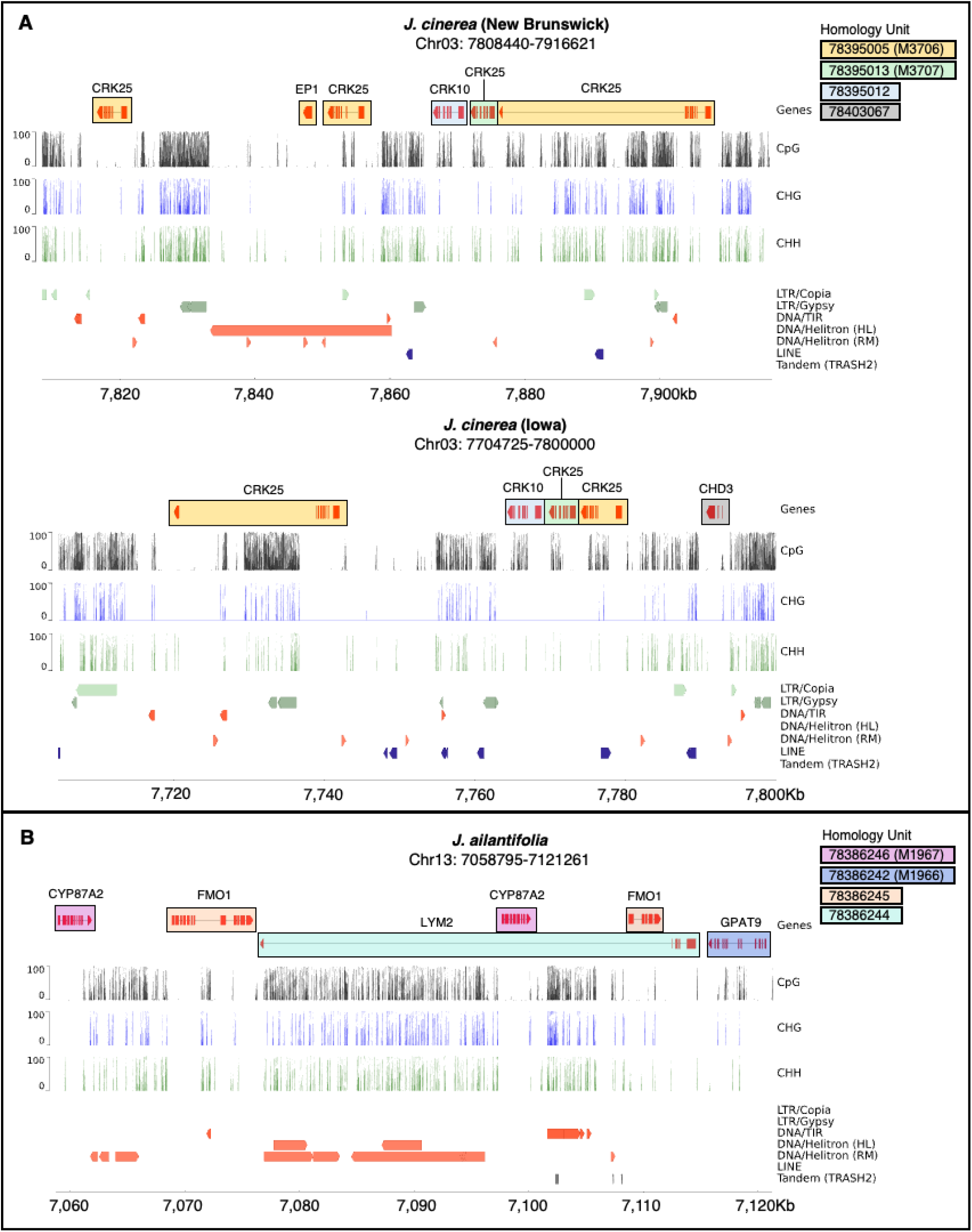
Landscape plots highlighting accession-specific gene organization. Tracks (top to bottom) show genes colored by homology unit, CPG, CHG, CHH methylation, and classification of *LTR/Copia*, *LTR/Gypsy*, *DNA/TIR*, *DNA/Helitron* (HL=HELIANO; RM=RepeatMasker), *LINE* and Tandem (TRASH2) repeat elements. **A)** Organization of cysteine-rich receptor-like kinase homology units 78395005 (yellow; *CRK25/EP1*), 78395013 (green; *CRK25*), and 78395012 (light blue; *CRK10*) in the *J. cinerea* New Brunswick (top; Chr03:7808440-7916621) and Iowa (bottom; Chr03:7704725-7800000) accessions, where one *DNA/Helitron* (HL) uniquely flanks 78395005 in New Brunswick. Chromatin-remodeling ATPase homology unit 78403067 (gray; *CHD3*) is uniquely identified in Iowa. **B)** Organization of cytochrome P450 associated unit 78386246 (pink; *CYP87A2*), glycerol-3-phosphate acyltransferase unit 78386242 (dark blue; *GPAT9*), flavin-containing monooxygenase unit 78386245 (orange; *FMO1*), and GPI-anchored protein 2 unit 78386244 (teal; *LYM2*) in *J. ailantifolia* (Chr13:7058795-7121261).

### Whole genome duplication across six Juglans genomes

Whole genome duplication was assessed across six *Juglans* genomes (*J. regia*, *J. ailantifolia*, *J. nigra*, *J. mandshurica*, and both *J. cinerea* accessions), with *Morella rubra* as an outgroup control. The within-genome comparison of *M. rubra* recovered no recent duplication, with Ks peaks at 1.01 and 1.99 consistent with the triplication shared among all eudicots and a further peak at 4.14 in the saturated range (Figure S7).

For the shared duplication history of the *Juglans* genomes, a mixed Ks distribution combining pairwise paralog and ortholog comparisons with anchor pair values was best fit by an exponential plus lognormal mixture model, which returned four components: a narrow peak near 0.3, a second at 0.47, and older components at 1.49 and 4.06 (Figure 7A). Anchor pairs support two of the four, at approximately 0.28 to 0.30 and 1.46 to 1.49 (Figure S8; Figure S9). These support interpreting the component near 0.3 as the Juglandaceae-specific Juglandoid WGD and the component near 1.5 as the shared eudicot γ-WGD (Bowers et al., 2003; Ding et al., 2023; X. Li et al., 2022).The remaining two components are not interpreted as duplication events. The component at 0.47 lies immediately above the Juglandoid peak and has no corresponding mode in the anchor pair distribution, which is expected where a mixture model fits a second lognormal to the right tail of a single peak, since among-gene variation in substitution rate makes the distribution generated by one duplication right-skewed. The component at 4.06 lies beyond the range over which Ks can be estimated reliably, because synonymous sites approach saturation above Ks of approximately 2, so its fitted mode is a lower bound.

**Figure 7.**
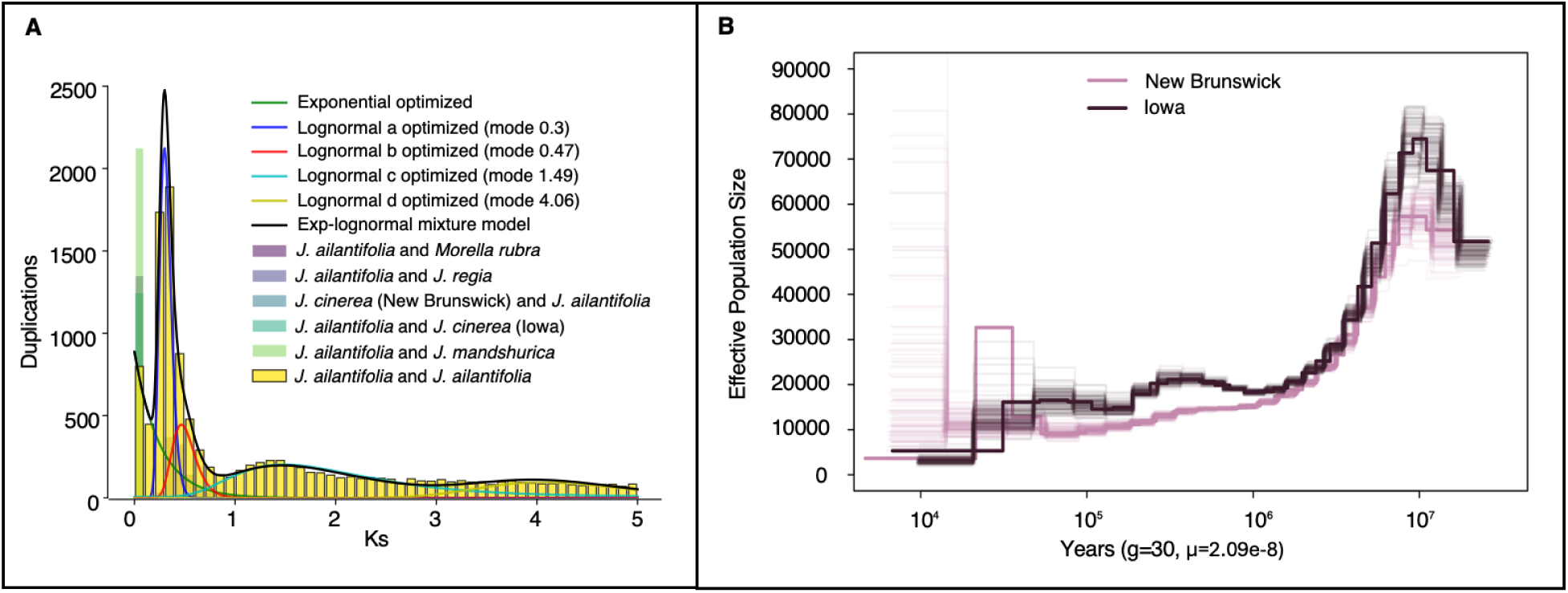
Gene duplication and population history in *Juglans.* **A)** Ks age distribution of the paranome of Juglandaceae, with *Juglans ailantifolia* as the focal species. The histogram shows the frequency of duplicated gene pairs across the x-axis, with the y-axis representing the duplication counts. The yellow bars represent the mixed Ks distribution of *J. ailantifolia*, while the other colored stacked bars show the contributing pairwise comparisons: *J. ailantifolia* and *Morella rubra* in purple, *J. ailantifolia* and *J.regia* in lavender, *J. cinerea* (N.B.) and *J. ailantifolia* in blue-grey, *J. ailantifolia* and *J. cinerea* (Iowa) in teal, and *J. ailantifolia* and *J. mandshurica* in light green. Overlaid models fits include an optimized exponential background model (green), four lognormal components with modes at 0.3, 0.47, 1.49, and 4.09 (blue, red, cyan, yellow) and the full exponential-lognormal mixture model in black. **B)** PSMC coalescence plot showing effective population size (Ne) of the *J. cinerea* Iowa (dark) and New Brunswick (light) accessions over time (10⁷ to 10⁴ years before present) using mutation rate 2.09e-8 and a generation time of 30 years.

### Demographic history of Juglans cinerea

SNP mapping and filtering produced 1,520,977 biallelic sites in the Iowa accession and 991,856 in the genetically distinct N.B. accession. Heterozygosity, calculated as the number of SNPs divided by the number of bases analyzed (491,720,421), was 0.309% in Iowa and 0.202% in N.B., within the range reported by Bai et al. (2018). Effective population size (*Ne*) at 10 Mya was largest in the Iowa accession (∼70,000) and smallest in the N.B. accession (∼60,000). Both then declined sharply to below 10,000, with the Iowa accession showing greater recovery around 1 Mya and 100 Kya (Figure 7B). Because PSMC has limited resolution at recent timescales, the ∼32 Kya rise in the N.B. accession is likely an artifact.

## DISCUSSION

### New Juglans assemblies extend the comparative framework

The first chromosome-scale reference for *J. ailantifolia* and a second *J. cinerea* accession from the primary range enable within-species and between-section comparison, and re-annotating *J. mandshurica* with the same pipeline removes annotation method as a source of variation. All three new or revised assemblies resolve into the expected 16 chromosomes and are comparable in completeness, contiguity and consensus statistics (Table 1; Figure 2). Re-scaffolding the N.B. reference with independently generated Hi-C reads preserves completeness while removing the structural bias toward *J. mandshurica*.

Chromosome-scale references are now available for ten *Juglans* species spanning all four sections, including Trachycaryon. The synteny-anchored analysis of the *J. ailantifolia* paranome resolves two duplication components with anchor-pair support (Figure 7A; Figure S8; Figure S9). The younger peak is interpreted as the Juglandoid duplication and the older as the eudicot triplication, consistent with recent phylogenomic work (X. Li et al., 2022; Ding et al., 2023).

### Section-level defense differences appear in gene presence and absence and in the copy number of five families

The two comparative frameworks answer different questions. CAFE asks where on the phylogeny a family changed size, using one genome per species and therefore representing *J. cinerea* by the N.B. accession alone (Figure 4). The pangenome asks how far a between-section difference exceeds that between two individuals of one species (or genus), and without a tree cannot distinguish common ancestry from convergence (Figure 5).

Both Cardiocaryon species show more than twice as many significantly changing terminal-node families as *J. cinerea*, whose signal is dominated by contraction across receptor and secondary metabolic families (Figure 4B; Table S17). Comparable contraction of surface receptor families was reported in *J. hindsii*, where the loss of FERONIA-like receptor kinases was proposed to underlie that species’ resistance to *Armillaria mellea* (Trouern-Trend et al., 2020). Receptor contraction is therefore uninformative about disease response on its own, appearing here in the susceptible species after being linked to resistance elsewhere in the genus. The directions here likewise do not track section membership: cysteine-rich receptor-like kinase 8 and the TMV resistance protein N-like family contract in both *J. cinerea* and *J. ailantifolia* while expanding in *J. mandshurica* (Table S17).

Six families differ between sections in copy number while remaining stable between the two *J. cinerea* accessions; five exceed the threshold and a sixth approaches it (Table S21). All favor Trachycaryon, and two modify pectin. Pectin is the substrate class most directly contested during bark colonization, since the cell wall is both the barrier a necrotroph must breach and a source of the fragments that trigger defense signaling (Miedes et al., 2014). The *Oc-j* genome also has an expanded complement of secreted CAZymes (Wu et al., 2020). A bias toward pectin-modifying capacity in the susceptible section is consistent with cell wall remodeling during infection and with constitutive defense.

Five defense families are enriched more than twofold among the section-specific units of both lineages (Table S21). Turnover in these families is shared by both lineages, and the balance of gain and loss explains why their summed copy number does not separate the sections. Wall associated kinases are the best characterized of the five in this genus (M. Li et al., 2022) and act as cell-wall-localized pattern-recognition receptors detecting oligogalacturonide damage signals (Stephens et al., 2022; Giovannetti and Genre, 2024). The asymmetric families divide by function: chitinases, lipoxygenases, NB-LRR receptors and germin-like proteins toward Cardiocaryon, and dehydrins, pathogenesis-related CRISP proteins, the CBF/DREB transcription factors and pectin-modifying enzymes toward Trachycaryon. Of the four Cardiocaryon-biased families, two act against fungi without receptor recognition: lipoxygenases initiate jasmonate synthesis (Bell et al., 1995), the signaling type most effective against necrotrophs (Macioszek et al., 2023), and germin-like proteins function collectively as a broad-spectrum resistance QTL (Manosalva et al., 2009). The NB-LRR receptors form a large, structurally variable family in *Juglans* (Chakraborty et al., 2016; H. Zhou et al., 2023), and chitinases are treated below. The strongest single enrichment is the dehydrin bias toward Trachycaryon, a cold response signal revisited below with the cold adaptation results.

Chitin defense expands in both Cardiocaryon lineages and shows the strongest Cardiocaryon bias in the pangenome, with cell-surface receptor signaling enriched among the same homology units (Figure 4B; Figure 5C; Figure 5E; Tables S17, S23, S25). Since chitin is the diagnostic component of the fungal cell wall, the more BCD-resistant Cardiocaryon may be better equipped for its perception and degradation. Terpenoid metabolism shows the opposite, with terpene cyclase expanding in *J. cinerea* and sesquiterpenoid biosynthesis and monoterpene synthase activities enriched among Trachycaryon-specific units. Terpene synthases diversify rapidly and generate lineage-specific profiles with both direct antimicrobial and indirect defense roles (Tholl, 2006).

### The Juglans sections differ in the composition of their defense complements

Cardiocaryon-specific gene content spans surface and intracellular immunity. Surface perception is represented by wall associated and cysteine-rich receptor-like kinases, and intracellular immunity by enrichment for the plant-type hypersensitive response together with three genes covering the known routes to it. Non-race-specific disease resistance 1 (NDR1) is required by coiled-coil NLRs and dispensable for TIR-domain receptors (Century et al., 1995; Aarts et al., 1998), senescence associated carboxylesterase 101 (SAG101) acts within the EDS1 complex that TIR-domain receptors require (Feys et al., 2005), and resistance to *Peronospora parasitica* 13 (RPP13) depends on neither (Aarts et al., 1998). Sampling more than one intracellular route is consistent with communication between the types of immunity (Ngou et al., 2021).

Chitin recognition is represented through lysin motif domain containing GPI-anchored protein 2 (LYM2), which restricts chitin-induced flux through plasmodesmata independently of CERK1-mediated immunity (Faulkner et al., 2013). This type of containment is directly relevant to *Oc-j*, which spreads through bark and girdles the stem progressively, so the rate at which a lesion is walled off may matter as much as the strength of initial recognition.

Salicylic acid distinguishes the sections by position within the pathway. Cardiocaryon is enriched for salicylate biosynthesis and its regulation, the isochorismate synthase route supplying most pathogen-induced salicylate (Wildermuth et al., 2001), while Trachycaryon is enriched for salicylate metabolism and carries a methyl esterase of the family that releases free salicylate from its methylated transport form (Vlot et al., 2008). One section is more focused on producing the signal and the other on processing it.

The juglone pathway supports a partial Cardiocaryon signal that does not generalize. Bark juglone may correlate with BCD resistance (Moore et al., 2015), yet the *Oc-j* genome hosts a large P450 complement likely involved in juglone detoxification (Wu et al., 2020). Gene family analysis recovers a CYP450 expansion at the ancestral Cardiocaryon node including the juglone pathway P450 (X. Li et al., 2022; Table S17), consistent with P450 diversification with lineage-specific metabolites (Mizutani and Ohta, 2010). In the pangenome, the largest P450 arrays fall in the phenylpropanoid branch, and the enrichment of phenylalanine catabolism among Trachycaryon-specific homology units supports this association. Anthracnose response in *Juglans* also implicates phenylpropanoid metabolism (Pollegioni et al., 2012), so this is likely not specific to *Oc-j* response.

### The largest copy number contrast is associated with a single Juglans cinerea accession

The largest copy number difference recovered separates the two *J. cinerea* individuals. The senescence associated protease gene SAG12 is present at high copy number in the Iowa accession and both Cardiocaryon genomes and at a single copy in N.B. (Table S20). It encodes a papain-like cysteine protease driving senescence associated proteolysis and nitrogen remobilization from aging leaves (Noh and Amasino, 1999; James et al., 2018). Gene family analysis recovers the same contrast as the largest contraction at the *J. cinerea* terminal node (Figure 4B; Table S17). Whether the array contributes to defense is uncertain. Senescence associated proteolysis intersects the programmed cell death that necrotrophs exploit, and *Oc-j* kills host tissue as it advances, yet SAG12 knockouts in *A. thaliana* show little phenotypic response (James et al., 2018). The link between senescence and immunity is better established at SAG101, also restricted to Cardiocaryon, and with a defined role in TIR-domain receptor signaling.

Leaf senescence regulation is also enriched in the Cardiocaryon-specific set, which is consistent with a phenological difference between the sections. *Juglans ailantifolia* leafs out earlier and is less cold-hardy than *J. cinerea*, and controlled trials show that buartnut hybrids are intermediate, with some hybrids possibly exceeding the cold tolerance of *J. cinerea* at its northern range limits (Brennan et al., 2021).

### Helitron associated structural variation is accession-specific

Helitrons capture host gene fragments as they mobilize, generating copy number variation and relocating gene copies (Kapitonov and Jurka, 2001; Barro-Trastoy and Köhler, 2024), a process associated with loss of gene content collinearity between maize inbred lines (Lai et al., 2005; Lynch et al., 2015). With the exception of dehydrins, the defense families that distinguish the sections here include tandemly arrayed members (Table S22).

Both *J. cinerea* accessions have the most complete Helitron elements, including those adjacent to protein-coding genes (Table S12). Across all four genomes the majority of elements retain the replication initiator and helicase machinery of autonomous Helitrons (Dong et al., 2011), and the resistant species contain the highest proportion of autonomous elements among Helitron-flanking genes. In keeping with their capacity to capture and relocate host gene fragments and to generate copy number and presence/absence variation, the Helitrons recovered here flank stress- and defense associated genes in an accession-specific pattern (Lai et al., 2005; Yang and Bennetzen, 2009; Grabundžija et al., 2016; Barro-Trastoy and Köhler, 2024). The antifungal CRK25/EP1 gene array in N.B. and an added copy of the systemic acquired resistance regulator FMO1 (Chen et al., 2018) in *J. ailantifolia* are associated with Helitrons of contrasting autonomy and methylation (non-autonomous and hypo-methylated versus autonomous and hyper-methylated), which may impose distinct epigenetic control over the captured genes (Barro-Trastoy and Köhler, 2024). We interpret these associations as accession-specific structural variation that supports defense gene turnover within a TE-mediated host-pathogen arms race (Fouché et al., 2022).

### Demographic histories separate the Juglans cinerea accessions, and cold adaptation differs in effectors while regulators are conserved

Coalescent trajectories were reconstructed from a single diploid genome per accession, which confines interpretation to the window from roughly 100 Kya into the late Miocene and leaves little power at recent timescales, where unmodeled population structure can generate spurious peaks (Li and Durbin, 2011; Hilgers et al., 2025). The two accessions share a comparable ancestral size and a common decline, after which their histories separate, with the Iowa accession recovering twice while the northern accession remains low (Figure 7B). Both read sets were mapped to the N.B. assembly, so reference bias should reduce the Iowa heterozygosity estimate and the observed direction is conservative. The admixed nuclear and plastid ancestry of *J. cinerea* (Aradhya et al., 2007; Mu et al., 2020) may also increase the inferred ancestral size under the panmixia assumption, and a prolonged period at low effective size is equally consistent with sustained isolation of a population as with a distinct refugial history.

Range-wide microsatellite data support this contrast. In the first range-wide survey, the N.B. populations show M-ratio evidence of historical bottlenecks, and the most northeastern population has the lowest allelic richness and expected heterozygosity of those surveyed (Hoban et al., 2010). Recent decline is unlikely to explain the pattern, since a seven-year reassessment identified reduced allelic richness in every breeding population and minimal changes in expected heterozygosity (van der Meer et al., 2026). Whether the northern population is a glacial refugium remains unresolved, though the organellar evidence favors an origin predating the last glacial advance. Laricchia et al. (2015) interpreted the low plastid diversity and divergent nuclear genotypes as preglacial differentiation. Resampling with additional northern accessions confirmed that the population is distinct and, together with fossil pollen and habitat-suitability estimates, supported a cryptic refugium while retaining a postglacial bottleneck as an alternative explanation (Schumacher et al., 2022).

The two *J. cinerea* accessions have comparable numbers of unique homology units in the pangenome, but those units differ in function (Table S26). The N.B. accession is enriched for calcium transport, regulation of salt stress responses and raffinose family oligosaccharide biosynthesis, while the Iowa accession is enriched for organic acid and amino acid transport and mitochondrial energy metabolism. Calcium influx is the first signal in cold perception and acts upstream of the CBF transcription factors (P. Yuan et al., 2018), and raffinose family oligosaccharides are cryoprotective osmolytes that accumulate during acclimation (ElSayed et al., 2014).

Cold adaptation in *Juglans* differs in downstream effectors, while the upstream regulators are conserved across sections. Partial CBF loss has been reported in species from mild winter climates, with more cold-adapted species keeping intact copies (Ebrahimi et al., 2020; Ebrahimi et al., 2025). In our four accessions the named cold-response regulators were invariant in copy number, whether at one copy (CBF1, CBF4, DREB1A, DREB2A, HOS10, HOS15) or two (CBF2, ICE1, HOS1), with the exception of a calmodulin binding transcription activator unit found in *J. mandshurica* (Table S20). The Trachycaryon bias for the CBF/DREB regulon is supported by a single section-specific homology unit. Controlled freezing trials report higher cold hardiness gene expression and less freezing injury in *J. cinerea* than in either East Asian species (Ebrahimi et al., 2020). The gene content differences appear further along the pathway. The dehydrin bias, the largest section-specific enrichment recovered, is associated with a contiguous block of three dehydrins and two LOB-domain proteins present in both *J. cinerea* accessions and absent from both Cardiocaryon genomes (Table S20). Within *J. cinerea*, the N.B. accession is additionally enriched for raffinose family oligosaccharide biosynthesis, a cryoprotective osmolyte pathway (ElSayed et al., 2014; Table S26).

Designatable unit recognition currently relies on geographic disjunction and neutral marker differentiation. The accession-specific content reported here spans both the perception and the effector pathways of freezing tolerance, though direct evidence of heritable variation for adaptive traits is still lacking and requires population level assessment (van der Meer et al., 2026). Prioritizing the northern population complements the case for ex situ collection in the south, where populations retain higher heterozygosity and more species-wide diversity (Hoban et al., 2010).

### Butternut canker tolerance is likely polygenic

Plant host defense ranges from single dominant R genes, such as the *Cr* loci conferring white pine blister rust resistance (J. Liu et al., 2022), to the quantitative resistance characteristic of chestnut blight (Westbrook et al., 2020). Both architectures occur within *Juglans*: resistance to crown gall and *Phytophthora* concentrates at a single major QTL in *J. microcarpa* × *J. regia* rootstocks (Saxe et al., 2025), while defense variation in the *J. nigra* pangenome is associated with numerous genes (H. Zhou et al., 2023). Butternut canker tolerance is likely to be polygenic, given that observed tolerance is rare, mostly confined to hybrids, and does not segregate as a few loci of large effect (Brennan et al., 2020; Conrad et al., 2026).

Resolving many loci of small effect relies on dense marker coverage and large mapping populations, which can be challenging in a long-lived outcrossing tree, especially in the absence of well resolved resistance candidate genes. Polygenic architecture does not preclude genetic gain, however. Field screening of a range-wide collection recovered high heritability of disease susceptibility at sites with sufficient disease pressure, with breeding values estimated for open pollinated families (Conrad et al., 2026). As such, selection can proceed on breeding values without resolving individual loci, and the chromosome-scale references reported here can supply the marker framework for this approach.

## Data Availability

The *J. cinerea* (Iowa) and *J. ailantifolia* ONT reads, genome assembly (pending for *J. cinerea*), and annotation (pending for *J. cinerea* and *J. ailantifolia*) are available through NCBI under Bioproject number PRJNA1151036 and PRJNA1055594 respectively.

## Acknowledgments

We thank the Institute for Systems Genomics Center for Genome Innovation at the University of Connecticut for molecular resources and support. We would also like to thank the Institute for Systems Genomics Computational Biology Core for access to HPC resources. Additional thanks to Kim Shearer for providing leaf tissue from Morton Arboretum.

## Supporting Information

**Figure S1.** Hi-C contact maps of *J. cinerea* (Iowa and N.B.) and *J. ailantifolia* assemblies distinguishing 16 chromosomes.

**Figure S2.** CpG, CHG, and CHH methylation patterns 5Kb up- and downstream from start and end coordinates of repeats: *LTR/Gypsy*, *LTR/Copia*, *DNA*/*TIR*, and *LINE*.

**Figure S3.** UpsetR plot representing total gene families shared by different intersections of 12 species in the Juglandaceae family.

**Figure S4.** Genespace synteny plot derived from OrthoFinder run.

**Figure S5.** Number of COGs functionally assigned to homology nodes within the pangenome by core , accessory and unique classes.

**Figure S6.** Log2 of protein lengths within core, accessory and unique homology nodes of pangenome.

**Figure S7.** K_s_ distribution for *Morella rubra* lacks a clear lineage-specific WGD peak. The histogram shows the weighted whole-paranome K_s_ distribution with fitted exponential and lognormal mixture components.

**Figure S8.** K_s_ age distribution of the paranome of Juglandaceae, with *Juglans ailantifolia* as the focus species.

**Figure S9.** Model selection for *Juglans ailantifolia* K_s_ age distribution. AIC and BIC both decrease from 1 to 2 components, indicating that a two-component model fits the data better.

**Table S1.** RNA libraries used to generate structural annotations.

**Table S2.** Public datasets downloaded for comparative analyses with genomes generated from this study.

**Table S3.** Oxford Nanopore sequencing data general summary before and after contaminants were removed.

**Table S4.** Genome size estimates from GOAT and kmer-freq, and final chromosome scale assembly size.

**Table S5.** Contaminants detected and removed from raw ONT reads with Centrifuge.

**Table S6.** Genome assembly metrics at each stage.

**Table S7.** FASTQC Summary of paired-end HiC reads.

**Table S8.** MassARRAY species status for *J. cinerea* and *J. ailantifolia* accessions.

**Table S9.** Final genome annotations produced by EASEL.

**Table S10.** Repeat landscapes from RepeatMasker.

**Table S11.** Weighted average Kimura divergence for each repeat family.

**Table S12.** Additional repeat classification with TRASH2 and HELIANO. **Table S13.** Transcripts surrounded by Helitrons output by HELIANO (+/- 5kb).

**Table S14.** OrthoFinder summary statistics.

**Table S15.** OrthoFinder N0.tsv output with EggNOG functional characterization from longest gene in gene family.

**Table S16.** OrthoFinder protein IDs per HOG.

**Table S17.** Rapidly expanding and contracting gene families output by CAFE (p<0.01).

**Table S18.** Pantools pangenome metrics.

**Table S19.** Transcript-level annotation, mapped to merged homology units.

**Table S20.** Copy number per merged homology unit across the four accessions.

**Table S21.** Gene family copy number and section-specific presence and absence.

**Table S22.** Tandemly arrayed genes by accession.

**Table S23.** Plant RefSeq functional annotation for transcripts in section-specific homology units.

**Table S24.** Araport11 functional annotation for transcripts in section-specific homology units.

**Table S25.** GO enrichment (BP and MF) of section Cardiocaryon and Trachycaryon-specific genes in pangenome (p<0.1).

**Table S26.** GO enrichment (BP) of Iowa and New Brunswick (N.B.) J. cinerea unique homology nodes in pangenome (p<0.1).

