## Supplementary material for "Chromosome-scale reference genomes for two walnuts: *Juglans ailantifolia* and *Juglans cinerea* implicate gene presence and absence in fungal defense divergence": Figure S1

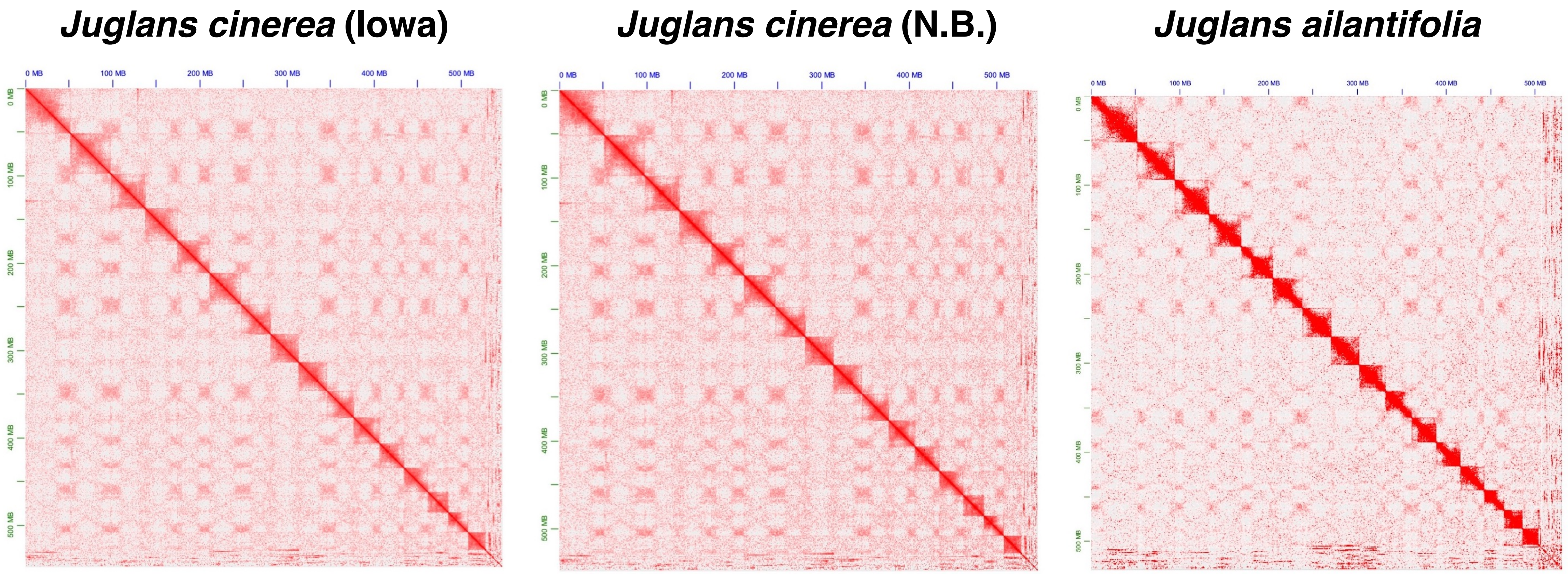


Figure S1. Hi-C contact maps of J. cinerea (Iowa and N.B.) and J. ailantifolia assemblies distinguishing 16 chromosomes. The color from light to dark shows an increase in interaction intensity.
