## Supplementary material for "Chromosome-scale reference genomes for two walnuts: *Juglans ailantifolia* and *Juglans cinerea* implicate gene presence and absence in fungal defense divergence": Figure S2

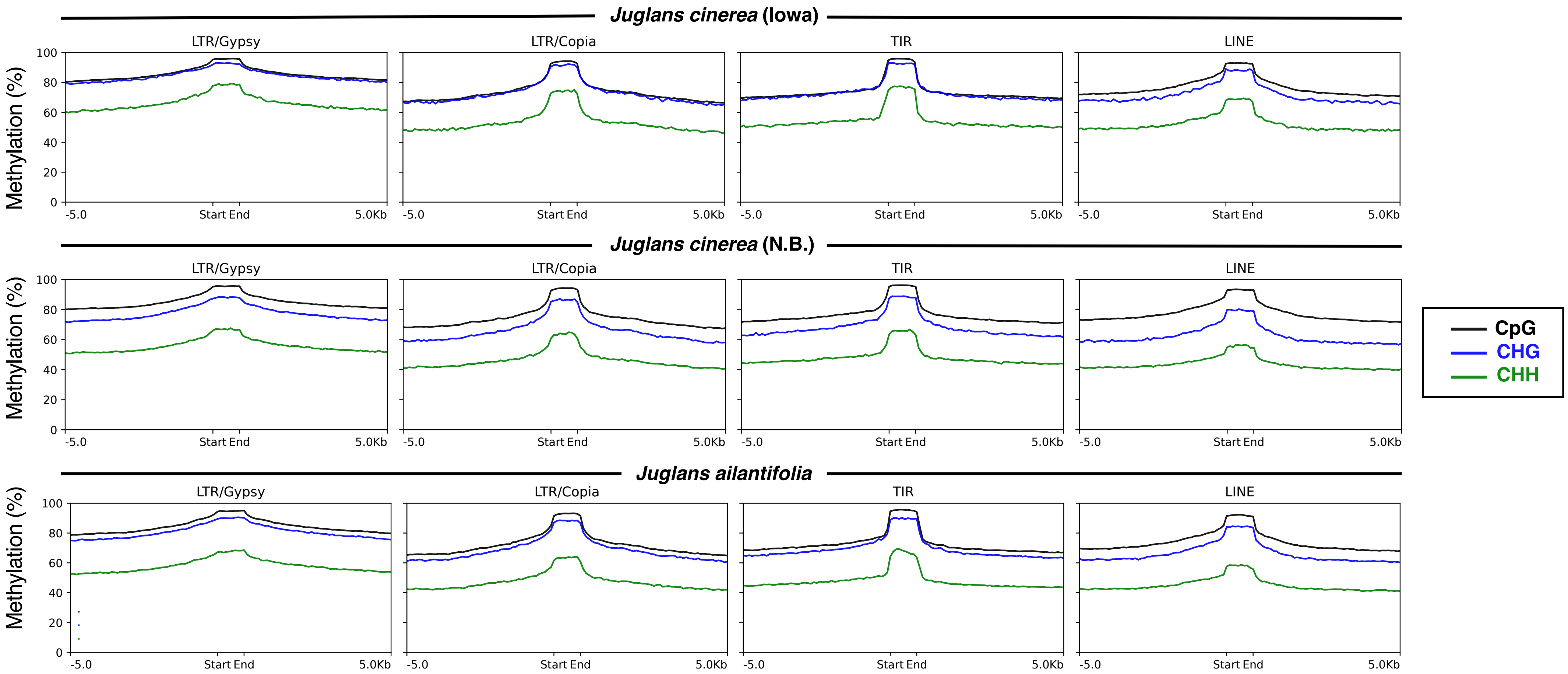


Figure S2. CpG (black), CHG (blue), and CHH (green) methylation patterns 5Kb up- and down-stream from start and end coordinates of repeats: LTR/Gypsy, LTR/Copia, DNA/TIR, and LINE.
