## Supplementary material for "Chromosome-scale reference genomes for two walnuts: *Juglans ailantifolia* and *Juglans cinerea* implicate gene presence and absence in fungal defense divergence": Figure S3

Figure S3. UpsetR plot representing total gene families shared by different intersections of 12 species in the Juglandaceae family. Juglans cinerea is represented by the N.B. accession.

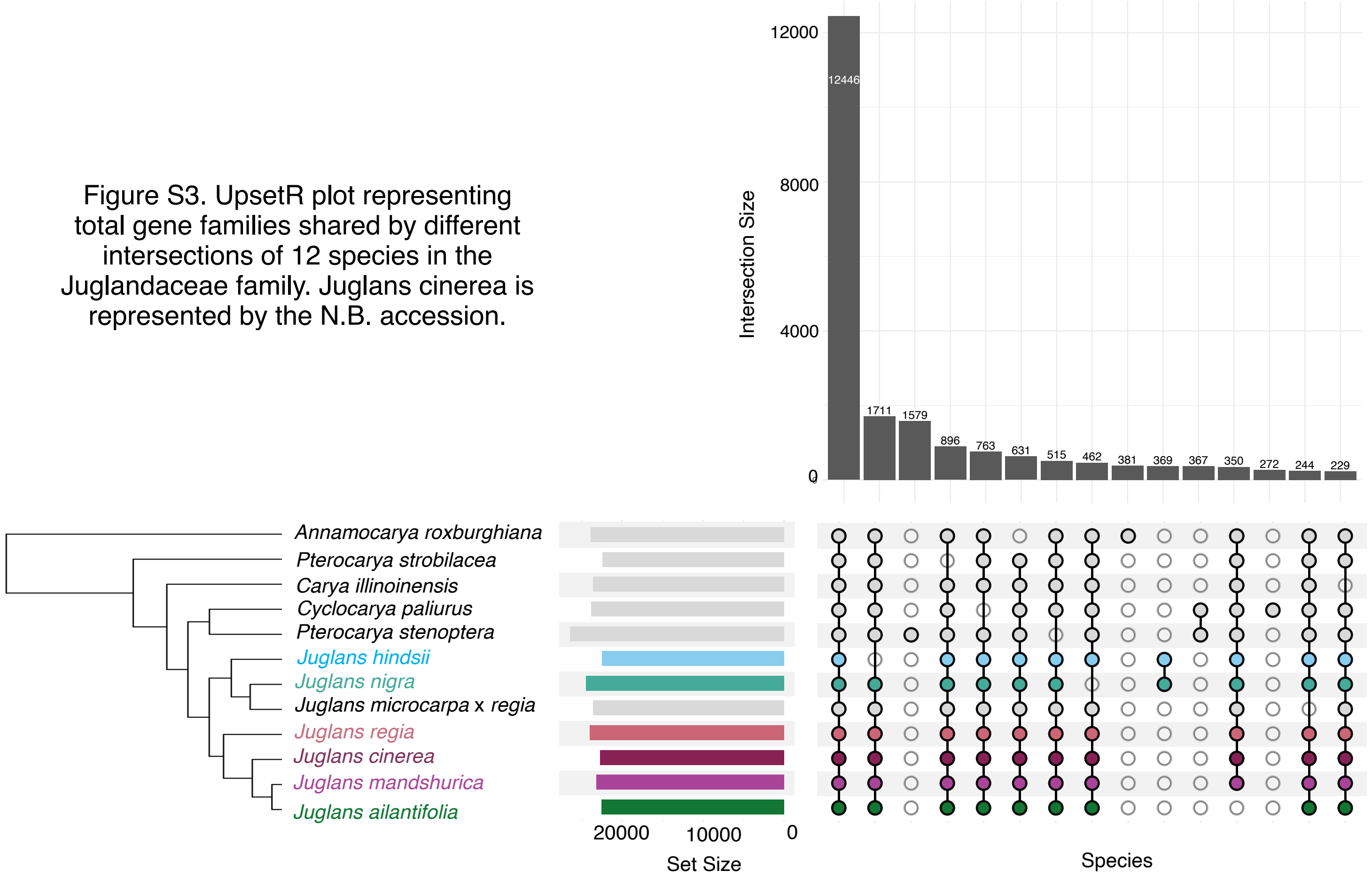
