## Supplementary material for "Chromosome-scale reference genomes for two walnuts: *Juglans ailantifolia* and *Juglans cinerea* implicate gene presence and absence in fungal defense divergence": Figure S4

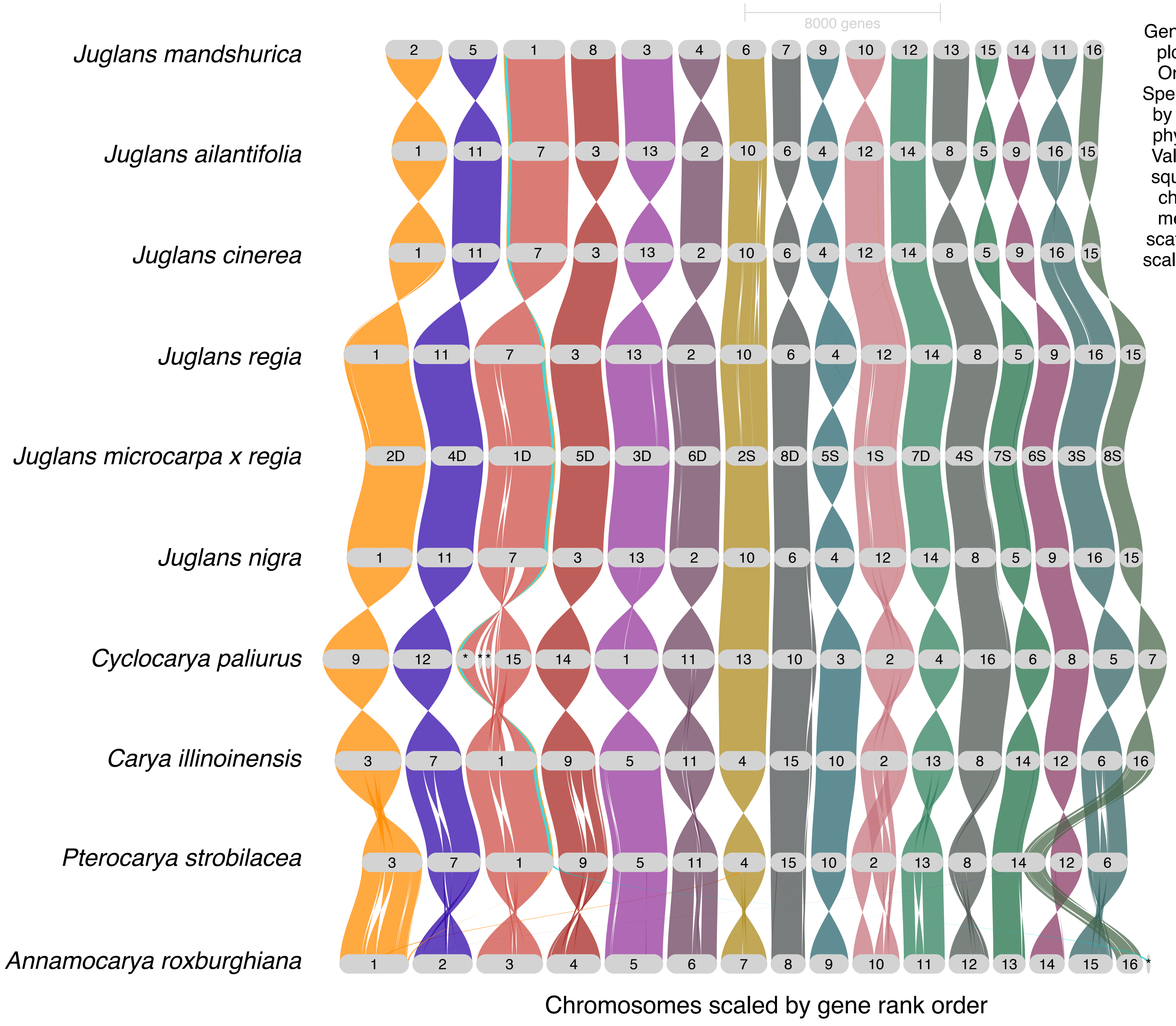


Figure S4. Genespace synteny plot derived from OrthoFinder run. Species are ordered by relatedness on phylogenetic tree. Values within grey squares represent chromosomes (* means unplaced scaffold), which are scaled by gene rank order.
