## Supplementary material for "Chromosome-scale reference genomes for two walnuts: *Juglans ailantifolia* and *Juglans cinerea* implicate gene presence and absence in fungal defense divergence": Figure S5

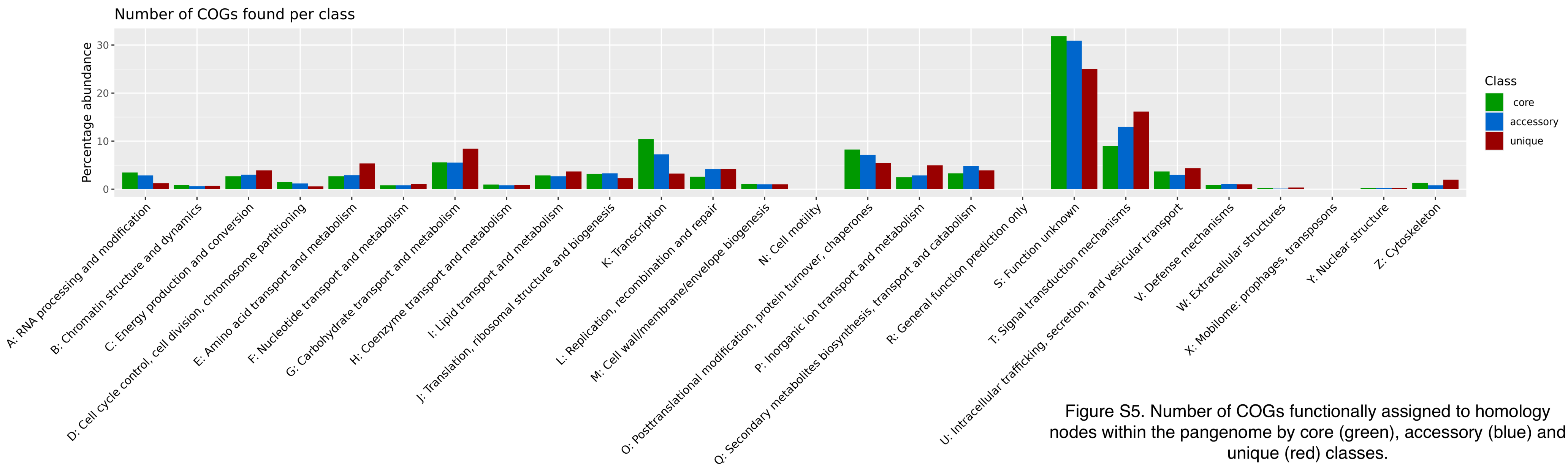


Figure S5. Number of COGs functionally assigned to homology nodes within the pangenome by core (green), accessory (blue) and unique (red) classes.
