## Supplementary material for "Chromosome-scale reference genomes for two walnuts: *Juglans ailantifolia* and *Juglans cinerea* implicate gene presence and absence in fungal defense divergence": Figure S6

Figure S6. Log2 of protein lengths within core, accessory and unique homology nodes of pangenome. Core compared to accessory and core compared to unique proteins show a significant difference (p=<0.0001). There is no significant difference between accessory and unique.

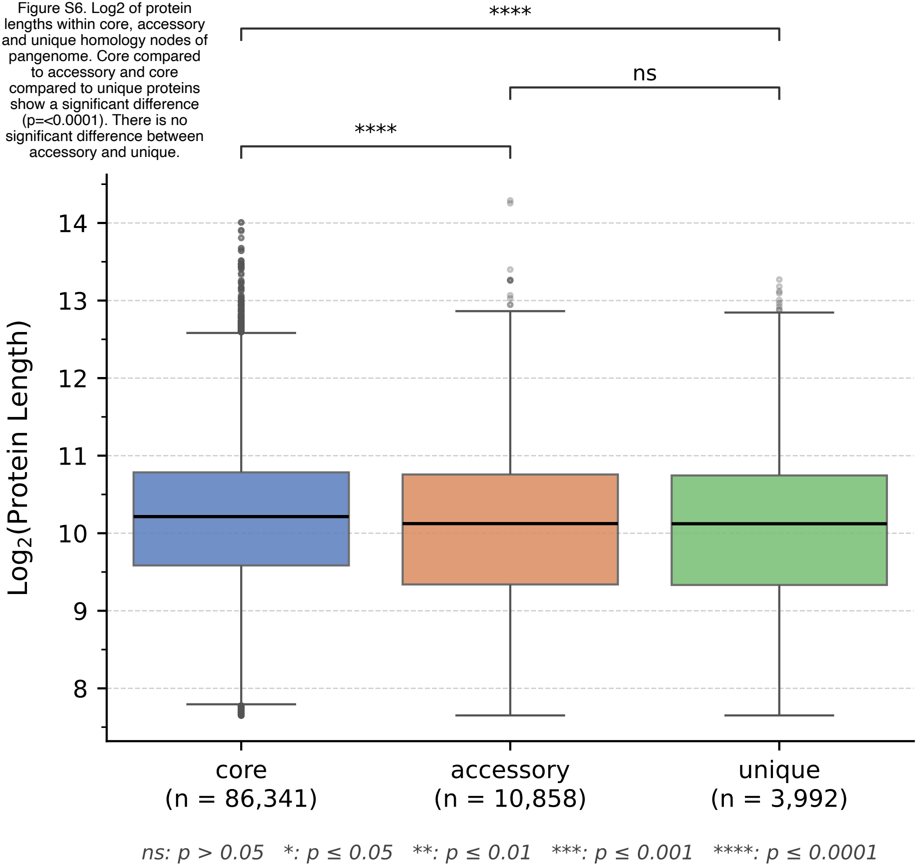
