## Supplementary material for "Chromosome-scale reference genomes for two walnuts: *Juglans ailantifolia* and *Juglans cinerea* implicate gene presence and absence in fungal defense divergence": Figure S7

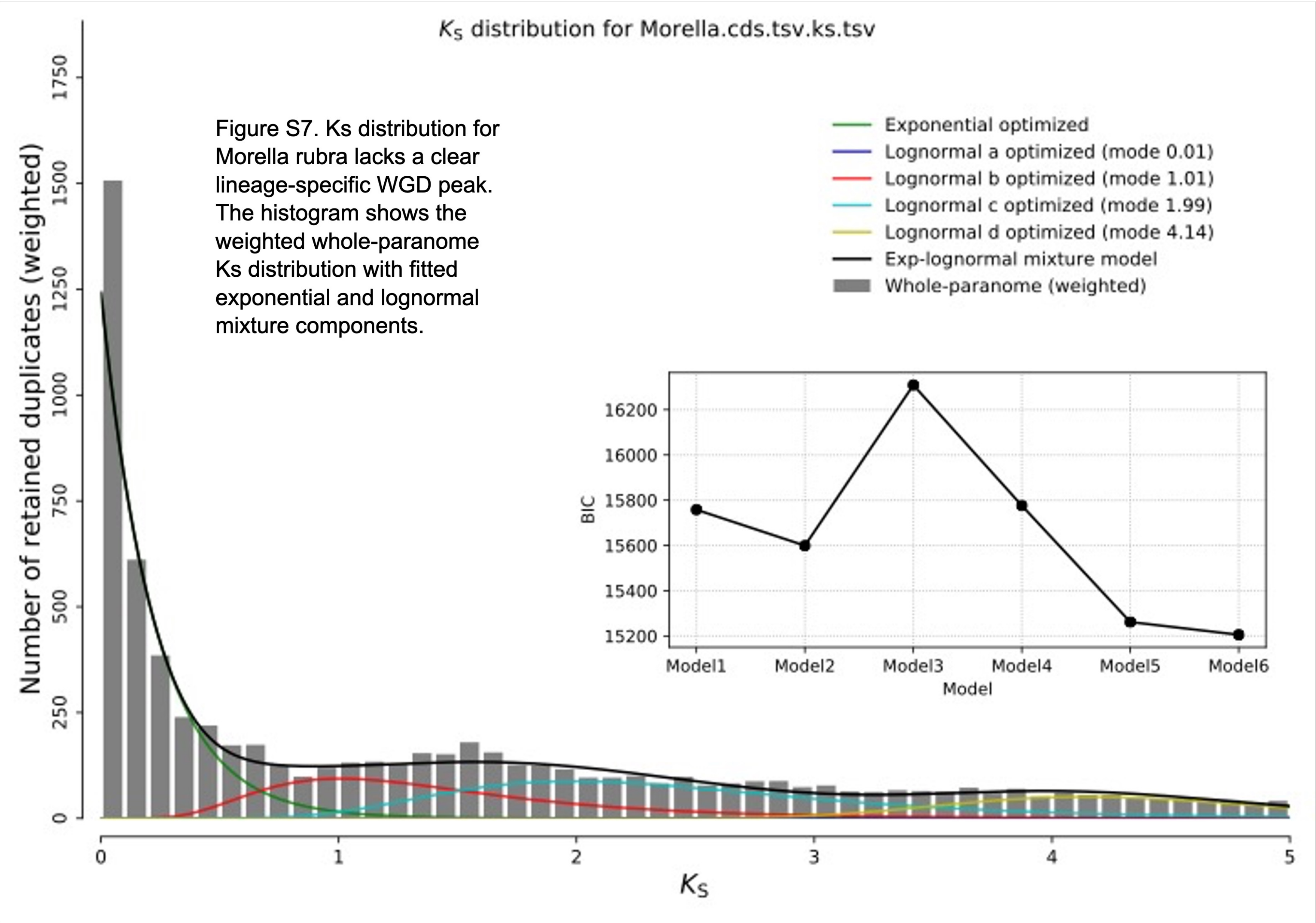


Figure S7. Ks distribution for Morella rubra lacks a clear lineage-specific WGD peak. The histogram shows the weighted whole-paranome Ks distribution with fitted exponential and lognormal mixture components.
