## Supplementary material for "Chromosome-scale reference genomes for two walnuts: *Juglans ailantifolia* and *Juglans cinerea* implicate gene presence and absence in fungal defense divergence": Figure S8

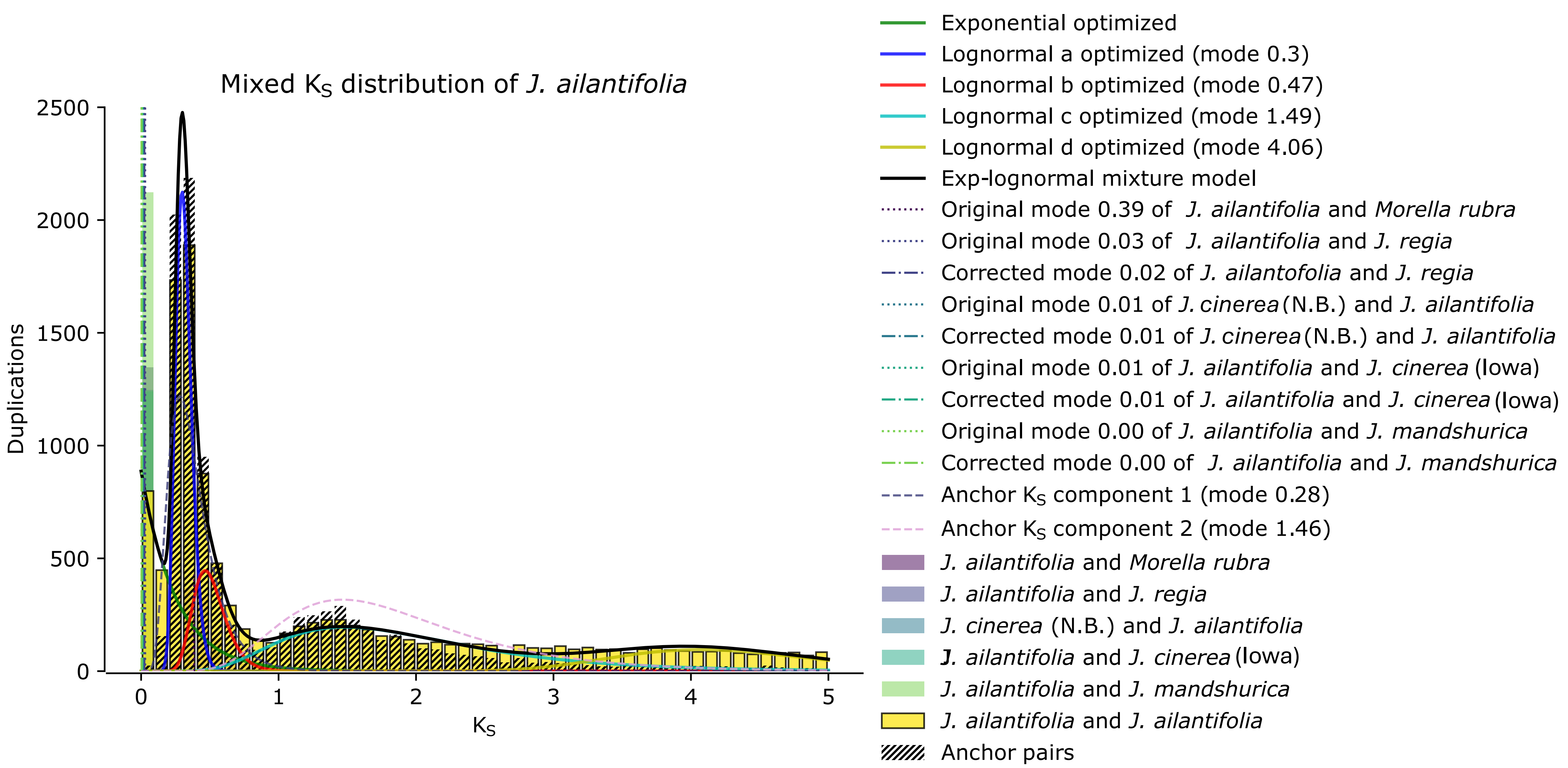


Figure S8. Ks age distribution of the paranome of Juglandaceae, with Juglans ailantifolia as the focus species. The solid lines show the inferred components of the mixture modeling analysis of the whole paranome, and the two anchor pair Ks distributions are denoted by the dashed lines. The corrected divergence times are denoted by the dash-dotted lines. The inlaid boxes show statistical significance in the choice of the model generated by wgd v2.
