## Supplementary material for "Chromosome-scale reference genomes for two walnuts: *Juglans ailantifolia* and *Juglans cinerea* implicate gene presence and absence in fungal defense divergence": Figure S9

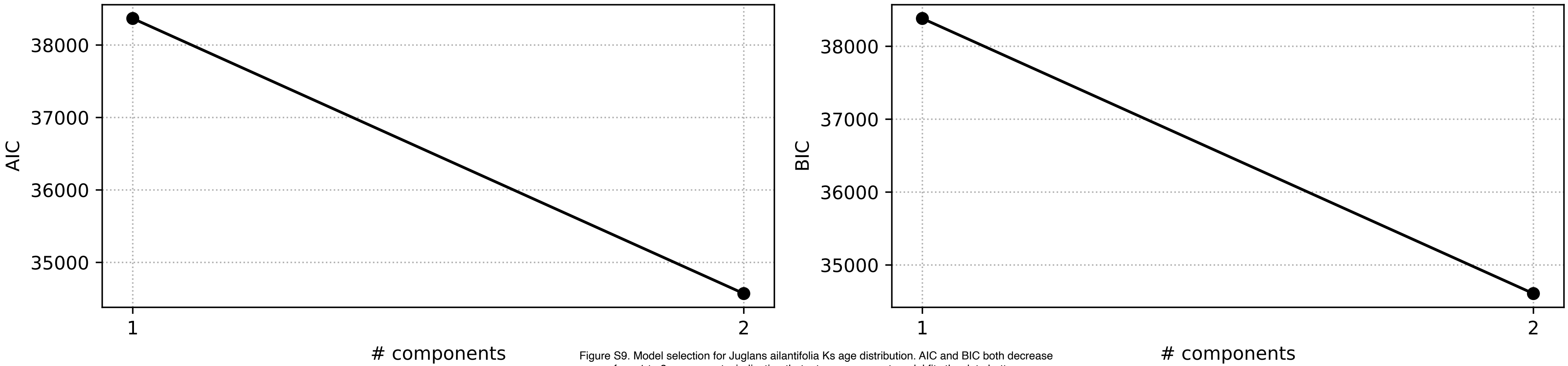


Figure S9. Model selection for Juglans ailantifolia Ks age distribution. AIC and BIC both decrease from 1 to 2 components, indicating that a two-component model fits the data better.
